# Nucleosomal DNA dynamics mediate Oct4 pioneer factor binding

**DOI:** 10.1101/634386

**Authors:** Jan Huertas, Caitlin M. MacCarthy, Hans Schöler, Vlad Cojocaru

## Abstract

Transcription factor (TF) proteins bind to DNA to regulate gene expression. Normally, accessibility to DNA is required for their function. However, in the nucleus the DNA is often inaccessible, wrapped around histone proteins in nucleosomes forming the chromatin. Pioneer TFs are thought to induce chromatin opening by recognizing their DNA binding sites on nucleosomes. For example, Oct4, a master regulator and inducer of stem cell pluripotency, binds to DNA in nucleosomes in a sequence specific manner. Here we reveal the structural dynamics of nucleosomes that mediate Oct4 binding. Nucleosome mobility and the amplitude of nucleosome motions such as breathing and twisting correlate with the number of Oct4 binding sites available. Moreover, the regions around the binding sites display higher local mobility. Probing different structures of Oct4-nucleosome complexes, we show that alternative configurations display stable protein-DNA interactions and are compatible with the DNA curvature and DNA-histone interactions.

---

Transcription factors (TFs) are proteins that directly or indirectly bind to DNA in order to promote transcribing genetic information. In most cases, accessibility to DNA is a prerequisite for TF binding. However, in the nucleus the DNA is packed into chromatin, making it often inaccessible for TF binding. The fundamental unit of chromatin is the nucleosome, which is formed by wrapping 147 DNA base pairs 1.65 times around a core of eight histones, DNA binding proteins consisting of a globular part and a disordered tail^1^. Packing DNA in this way leaves it partially occluded on the histone interface and by the second gyre of DNA.

Interestingly, a series of TFs are able to bind to closed, inactive chromatin, i.e. they recognize their binding sites even in the presence of histones. These factors, known as pioneer TFs, can help open chromatin, increase DNA accessibility, and support binding of other TFs. Among the TFs shown to bind to closed chromatin, several regulate transitions between cell fates^2^.

Particularly relevant for inducing pluripotency are Oct4, Sox2, and Klf4. These three TFs are known for their capacity to convert differentiated cells into pluripotent cells, a process known as reprogramming^3^, and have been characterized as pioneer TFs^4,5^. Structurally, Oct4 has two DNA binding domains (the POU specific domain, POU_S_ and the POU homeodomain, POU_HD_) that recognize an octamer sequence in free DNA^6^. In pluripotent cells, Oct4 cooperates with Sox2 to recognize composite motifs of free DNA^7^ via mechanisms involving structural dynamics that we previously characterized^8–10^. However, the interaction of these TFs with nucleosomes remains obscure. A recent survey of TF-nucleosome binding properties was performed for many different classes of TFs revealing how the nucleosome modifies the orientation and position of the TF binding compared to free DNA. Moreover, many TFs analyzed were capable of inducing nucleosome dissociation upon binding^11^. In the case of Oct4, the canonical configuration known from the structures of Oct4 bound to free DNA^6^ is incompatible with nucleosome binding because it involves binding of the POU_S_ and POU_HD_ on opposite sides of DNA. Instead, a partial motif recognition model was proposed, where only one of the domains is bound sequence specifically to the nucleosome^5^.

Although nucleosomes can assemble on just about any sequence, certain DNA sequences have structural features that lead to a higher affinity for the histone core proteins. For example, stiff DNA and polyA or T tracts are unfavorable for nucleosome formation but bendable DNA as well as DNA with AA/TT or GC dinucleotides in 10bp periodicity prove favorable^12^. Accordingly, DNA sequence also influences nucleosome stability and mobility^13^. However, the relationship between DNA sequence and nucleosome dynamics is poorly understood, especially when it comes to nuances in motion (e.g. local flexibility, compaction, breathing, twisting).

Despite the recent spike in pioneer factor studies, an atomic resolution structure of a TF-nucleosome complex still doesn’t exist. Moreover, the interplay between TF binding and sequence dependent nucleosome dynamics is not understood. This prevents rational design of pioneer TFs for improving efficiency and, perhaps, the direction of cell fate transitions. Therefore, it is of utmost importance to elucidate the features of nucleosomes that facilitate TF binding and to reveal how TF binding modulates nucleosome dynamics.

All-atom molecular dynamics (MD) simulations provide a powerful approach to study nucleosome dynamics. By solving Newton’s equations of motion, the motions and interactions of biomolecules at atomic resolution can be studied. Nucleosome motions have been simulated^14–16^, and motions such as the opening and closing of the linker DNA, the twisting involved in nucleosome repositioning^17^, or the behavior of the linker DNAs in the presence of linker histones^18^ have been accurately described.

Nevertheless, all these previous atomic resolution MD simulations^19^ were started from the most complete crystal structure of the nucleosome^20^, using a palindromic DNA sequence derived from human α-satellite DNA. Interestingly, coarse-grained MD simulations, in which several non-hydrogen atoms are grouped in beads have been performed with different DNA sequences^21–23^. Many of these studies used the Widom 601 artificial sequence^22,24^, an artificial sequence selected for its stability^25^ that is present in many crystal structures of the nucleosome^26,27^. Due to the reduced representation, the computational cost of these simulations is much lower. Therefore, questions that require longer and a higher number of simulations can be addressed. As a result, models of sequence-specific nucleosome dynamics, revealing the role of the sequence in nucleosome breathing and unwrapping^22,28^, or protein-mediated remodeling^29^ have been established. However, the descriptions obtained from coarse grained simulations is limited because important details regarding the interactions that modulate nucleosome dynamics are missing. For example, the hydrogen bonds driving the interaction of histone tails with the DNA grooves or the fine structural properties of DNA modulating TF binding could not be studied. Therefore, atomic resolution methods are indispensable for revealing the interplay between TF binding and nucleosome dynamics.

To study the Oct4-nucleosome interaction, we selected 2 native nucleosome sequences shown to be bound by Oct4 by Soufi et al. during reprogramming ^4^. The sequences were discovered by overlapping datasets of two types of sequencing experiments: the Oct4 binding sites from chromatin immunoprecipitation (ChIP) with the nucleosome positioning from micrococcal nuclease (MNase) digestion. We selected these sequences for their role in pluripotent cells, clarity of nucleosome boundaries, and number of TF binding sites. They are found in the enhancers of the *Esrrb* and *Lin28b* genes. Esrrb is a TF that regulates cellular pluripotency, whereas Lin28b is an RNA-binding protein, expressed in pluripotent stem cells, and involved in the regulation of glucose metabolism. The Esrrb sequence has just one obvious POU_S_ binding site, whereas Lin28b has at least two sites that Oct4 can interact with, one for the POU_S_ and one for the POU_HD_. The Oct4-Lin28b binding strength and position was established by electrophoretic mobility shift assays (EMSAs) and footprinting^4^.

We first compared the dynamics on the microsecond (µs) time scale of the two native nucleosomes with those of an artificial nucleosome with the Widom 601 sequence from atomic resolution MD simulations. We found differences in the mobility as well as in the amplitude of different types of motions such as nucleosomal breathing, twisting or enhanced fluctuations among the nucleosomes, that correlated with the number of TF binding sites. We validated these findings using temperature-induced nucleosome disassembly. Then, we built models of Oct4 bound to a mononucleosome and show that some non-canonical configurations but not all tested configurations of Oct4-nucleosome complexes were stable and not affected by DNA curvature or DNA-histone interactions. Our findings describe nucleosome dynamics that facilitate Oct4-nucleosome binding and provide a model for the structural basis of Oct4-nucleosome recognition.

## Results

### Nucleosomes with multiple transcription factor binding sites are more dynamic

We performed three, 1 µs MD simulations for each the Widom, Esrrb, and Lin28b containing nucleosomes. Given that 1 µs is insufficient for convergent sampling of the conformational space of the nucleosome, we combined three individual simulations of each nucleosome into one ensemble and performed all analyses on the three ensembles obtained.

To characterize the nucleosome mobility, we first fit the structured regions of the core histones from each snapshot on the starting model of the Widom nucleosome, and calculated the atomic fluctuations of the DNA residues (Figure 1). Then, we converted the fluctuations in β-factor analogues to crystallographic β-factors and normalized the β-factors by the mean β value of the core DNA (see Methods). We found that the most mobile regions are the linker DNAs (L-DNAs). Remarkably, their motions propagate inside the core nucleosomal DNA (the 147 bp that define the nucleosome particle) and are asymmetric. To distinguish between the two L-DNAs, we used the genomic 5’-3’ orientation and referred to them as 5’ L-DNA and 3’ L-DNA. For the Widom nucleosome, we defined the 5’ and 3’ ends, as the ends labelled with negative and positive residue numbers respectively in the structure with the PDB ID 3LZ0^26^. The 5’ L-DNA displayed higher mobility in regions located deeper in the core DNA. Moreover, we detected small enhanced mobility patches inside the core DNA, independent of the L-DNA mobility.

**Figure 1.**
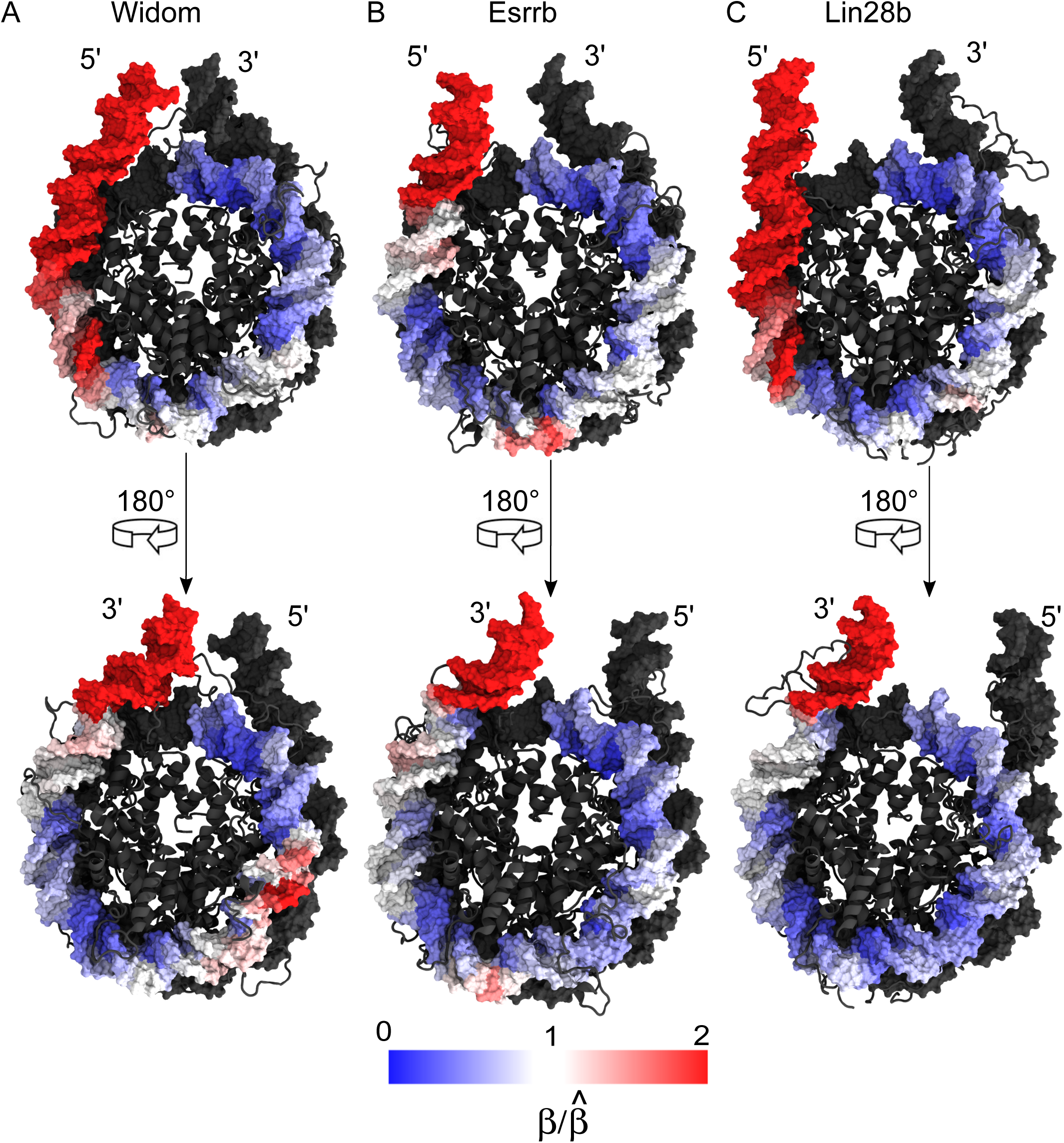
Propagation of asymmetric nucleosome flexibility. Normalized, relative atomic fluctuations of the non hydrogen DNA atoms calculated from the simulation ensembles of the 3 nucleosomes: **(A)** Widom, **(B)** Esrrb, **(C)** and Lin28b. First, the absolute atomic fluctuations were calculated, and the average (mass-weighted) per residue was then squared and weighted by 8/3π^2^ to obtain factors β analogues to crystallographic B-factors. The nucleosomes were colored by 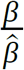, where 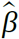 is the mean β of the core nucleosomal DNA (146 bp). The residues with fluctuations smaller and higher than the average fluctuations of the core DNA are in blue and red, respectively, with darkest red corresponding to 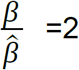 to avoid the masking of fluctuations by the very high values found in the linker DNA.

To study the differences in the mobility of the three nucleosomes, we calculated the distance root mean square deviation (dRMSD) of the DNA backbone (Fig 2A). The dRMSD measures the deviation of all pairs of internal distances from a reference structure and is independent of the initial fitting of the trajectory snapshots to the reference structure. The dRMSD values indicate how far different structures deviate from the reference, whereas their spread (width of the dRMSD histograms) is a measure of intrinsic mobility. We used the average structure of the DNA backbone from all simulations as the reference, ensuring that no reference dependent bias in the dRMSD values was introduced.

**Figure 2.**
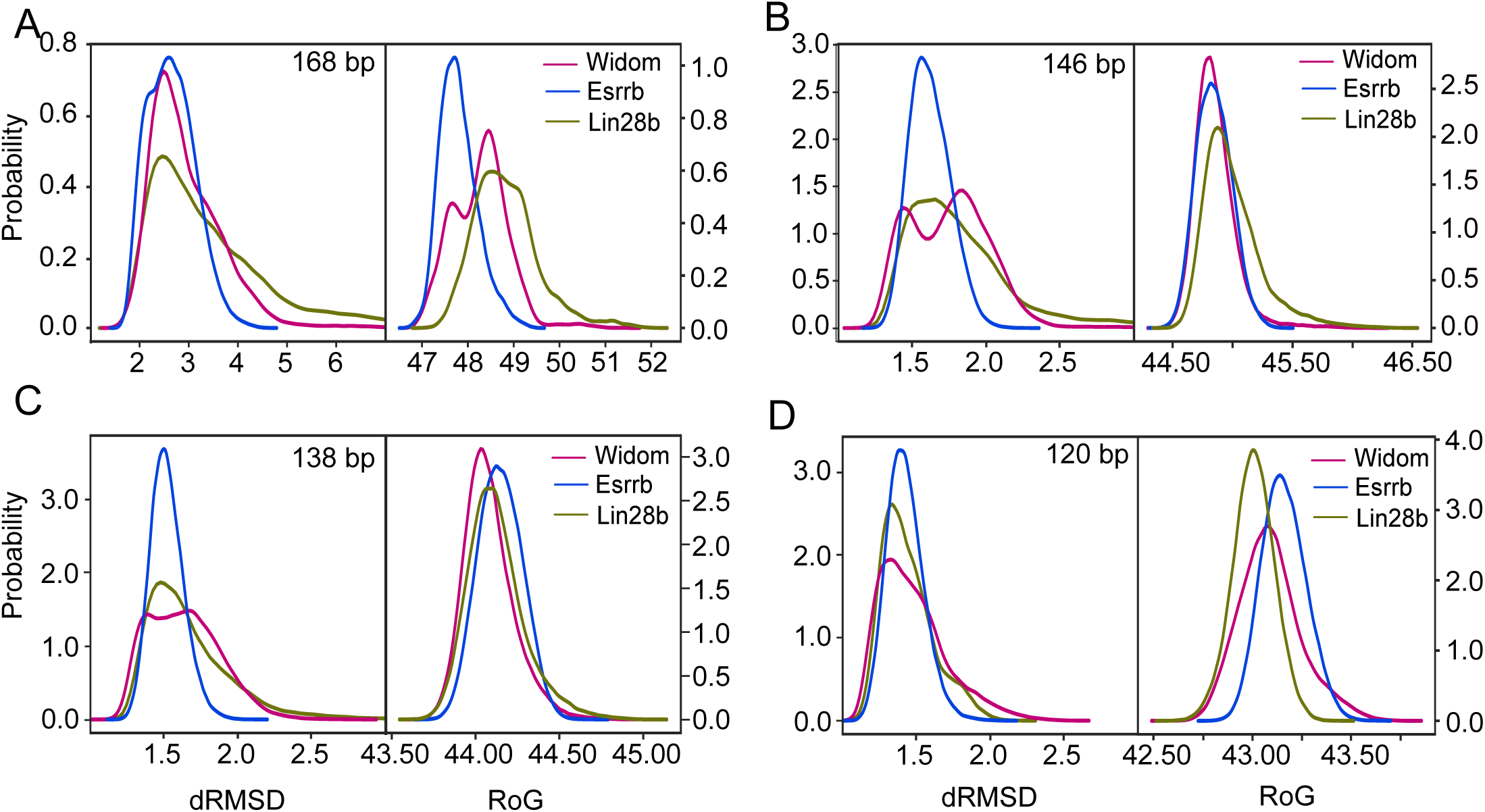
Sequence dependent nucleosome flexibility. Distributions of the root mean square deviation of interatomic distances in the DNA sugar-phosphate backbone (dRMSD, left) and radius of gyration (RoG, right) from the simulation ensembles of the 3 nucleosomes. The distributions were calculated for different regions of DNA: **(A)** The entire simulated DNA (168 bp), **(B)** The core nucleosomal DNA (146 bp) obtained by excluding 11 bp of linker DNA at each end, **(C)** Nucleosomal DNA obtained by excluding 15 bp of DNA at each end (138 bp), **(D)** Nucleosomal DNA corresponding to the complete inner DNA turn and half of the outer DNA turn obtained by excluding 23 bp of DNA at each end (120 bp).

To correlate the mobility with the compaction of the nucleosomes, we also calculated the radius of gyration of the DNA. Higher radius of gyration values reflect a decrease in nucleosome compaction. We found a hierarchy of mobility among the nucleosomes. The Lin28b, having the highest AT content (Table 1) and multiple Oct4 and other TF binding sites, was the most mobile. It was followed by the artificial Widom nucleosome with no Oct4 binding sites. The least mobile was the Esrrb that has a similar AT content as the Widom and contains a single Oct4 binding site (Table 1, Fig. 2A). A measurable population of the Lin28b nucleosome adopted conformations with high dRMSD values, indicating it undergoes larger conformational transitions compared to the other two. This hierarchy of mobility was correlated with the compaction of the nucleosomes, the Lin28b sampling less compact conformations (Fig 2A). The Esrrb nucleosome was the most compact (Fig 2A), whereas the Widom adopted two distinct populations, one as compact as the Esrrb and one as compact as the most compacted population of the Lin28b (Fig 2A).

**Table 1:** Overview of the simulated nucleosomes.

| **++Sequence** | **Origin** | **Oct4 Binding Sites** | **AT content (%)** | **Simulation time** | **Temperature of dissociation** |
| --- | --- | --- | --- | --- | --- |
| Widom | Artificial | 0 | 42.26 | 3 x 1 μs | 87.33 ± 0.58 |
| Esrrb | Natural | 1 | 45.24 | 3 x 1 μs | 86.00 ±1.73 |
| Lin28b | Natural | > 1 | 59.52 | 3 x 1 μs | 83.00 ±0.00* |

\*Standard deviation of 0.00 indicates that the maximum disassembly temperature measured was equal in the three independent experiments, even if the disassembly profile differed. See Fig S1 for more details.

Comparing the three independent simulations of each nucleosome, we observed that the Lin28b and the Widom sampled distinct regions of the conformational space in each simulation, whereas the conformations of the Esrrb are more similar among triplicates (Fig S2A). This confirmed that 1 µs is not sufficient for a converged sampling of the conformational space, and that merging the simulations into ensembles is necessary to obtain a more complete view of the nucleosome mobility. The intermediate mobility of the Widom nucleosome is due to motions mainly at the 5’ end that occurred in the second simulation, which include a short-lived large opening at the beginning of the simulation and a steady opening in the second half of the simulation (Fig. S2A, Fig. S3A). In addition, a partial opening at the 3’ end occurred after 750 ns in the third simulation (Fig. S2A, S3A). The higher mobility of the natural Lin28b nucleosome is not only rooted in the different sampling across different simulations but also in transitions occurring in the same simulation (Fig. S2A). The Lin28b undergoes large concerted opening-closing motions in the first two simulations and a smaller scale closing in the third simulation mostly at the 5’ end (Fig. S3A).

To validate the observed differences in motion between these nucleosomes, we performed an established thermal stability assay on reconstituted nucleosomes^30^. We found that the Widom had the highest mean temperature of dissociation in two of the three experiments performed (Table 1, Fig. S1). The Esrrb had an intermediate temperature of dissociation between the Widom and Lin28b. The large fluctuations of the measurements for this nucleosome suggest that its mobility varies. In one experiment, the Esrrb was more stable than Widom (Fig S1). Therefore, the low mobility of the Esrrb in the simulations reflects the dynamics within a subpopulation of its ensemble of conformations. The Lin28b had the lowest temperature of dissociation, requiring the least energy for disassembly. This confirmed our findings from the simulations that the Lin28b, a nucleosome with high AT content and multiple TF binding sites is the most mobile among the three nucleosomes studied.

### Differences in mobility between nucleosomes are not restricted to the linker DNA

To analyze the propagation of nucleosome mobility into the core region, we calculated the distributions of dRMSD for different fragments of DNA. First, we compared the mobility of the 146 bp DNA fragments that formally define the core nucleosomes. Interestingly, the core of the artificial Widom nucleosome displays two distinct peaks, indicating the sampling of two distinct conformational populations similar to the entire nucleosome (Fig. 2B). However, in the case of the core fragments, these two populations contain equally compact conformations as the radius of gyration has a very narrow, single peaked distribution (Fig 2B). The core of the Esrrb has a uniform, narrow dRMSD distribution reflecting its lower mobility (Fig 2B). Remarkably, the core of Lin28b displays a wide dRMSD histogram and adopts less compact conformations (Fig. 2B). Therefore, the increased mobility of the Lin28b nucleosome is propagated beyond the L-DNA arms and it involves decompaction of the core nucleosomal DNA. The most decompacted conformations were observed between 500 and 800 ns of the second simulation (Fig. S3B).

In general, the differences observed between the 146 bp core fragments are similar to those between the 138 bp DNA fragments, which are the largest fragments devoid of L-DNA motion propagation. However, the two conformational populations of the Widom nucleosome are less clearly separated (Fig. 2C), suggesting that these populations differ mainly in the L-DNAs and the adjacent 7 bp. The 138 bp fragment of Lin28b is still more mobile than the similar fragments of Widom and Esrrb (Fig 2C). This mobility was mainly apparent in the first two simulations (Fig. S3C). The main difference between the 146 bp and 138 bp fragments is in their compactness. The 138 bp fragments are very similar, and show no decompaction in the case of Lin28b (Fig. 2C, Fig. S3C). Therefore, the overall decompaction of Lin28b was due to conformational dynamics in the L-DNA arms and the adjacent 7 bp from the nucleosomal DNA.

Finally, the hierarchy of the mobility and compaction of the three nucleosomes changes when the inner 120 bp fragments were compared (Fig. 2D). The mobility of these fragments was lower and more similar between the nucleosomes as shown by the narrower dRMSD distributions (Fig. 2D). Interestingly, the Widom fragment shows the widest dRMSD distribution (Fig. 2D), suggesting that the inner part of this nucleosome is somewhat more mobile than the corresponding part of the two natural nucleosomes (Fig. 2D, Fig. S3D). In addition, the 120 bp fragment of Lin28b is the most compact (Fig. 2D, Fig. S3D), indicating that a higher overall mobility of the nucleosome is coupled with a less mobile and more compact region deep in the core around the dyad.

### The amplitude of nucleosome breathing motions depends on DNA sequence

To understand the origin of the sequence dependent nucleosome mobility and compaction, we characterized the different types of motions we observed. These were: breathing in which the L-DNAs open and close either parallel or perpendicular to the nucleosomal super-axis, changing its accessibility, DNA over-twisting, and local fluctuations.

To characterize the breathing motions, we defined two angles, γ1 and γ2, that describe the motions of the L-DNA on two orthogonal planes (see also Methods). γ1 describes a scissors like motion of the L-DNAs in which the dyad can be viewed as the point connecting the two blades, whereas γ2 describes a motion perpendicular to the plane of the blades. From the γ1-γ2 histograms (Fig 3C), we concluded that all three nucleosomes sampled different but overlapping regions of the conformational space in the three independent simulations (represented in magenta, gold, and cyan in Fig 3C). This indicates that although full convergence was not achieved in 1 µs, the most relevant parts of the conformational space were sampled. Further, we confirmed the asymmetry in the nucleosome mobility, the 5’ L-DNA sampling a larger area in the 2D histogram than the 3’ L-DNA, for all three nucleosomes. At last, we confirmed the higher mobility of Lin28b, especially at the 5’ end (Fig 3C, up right) and the lower mobility of Esrrb (Fig. 3C, middle column).

**Figure 3.**
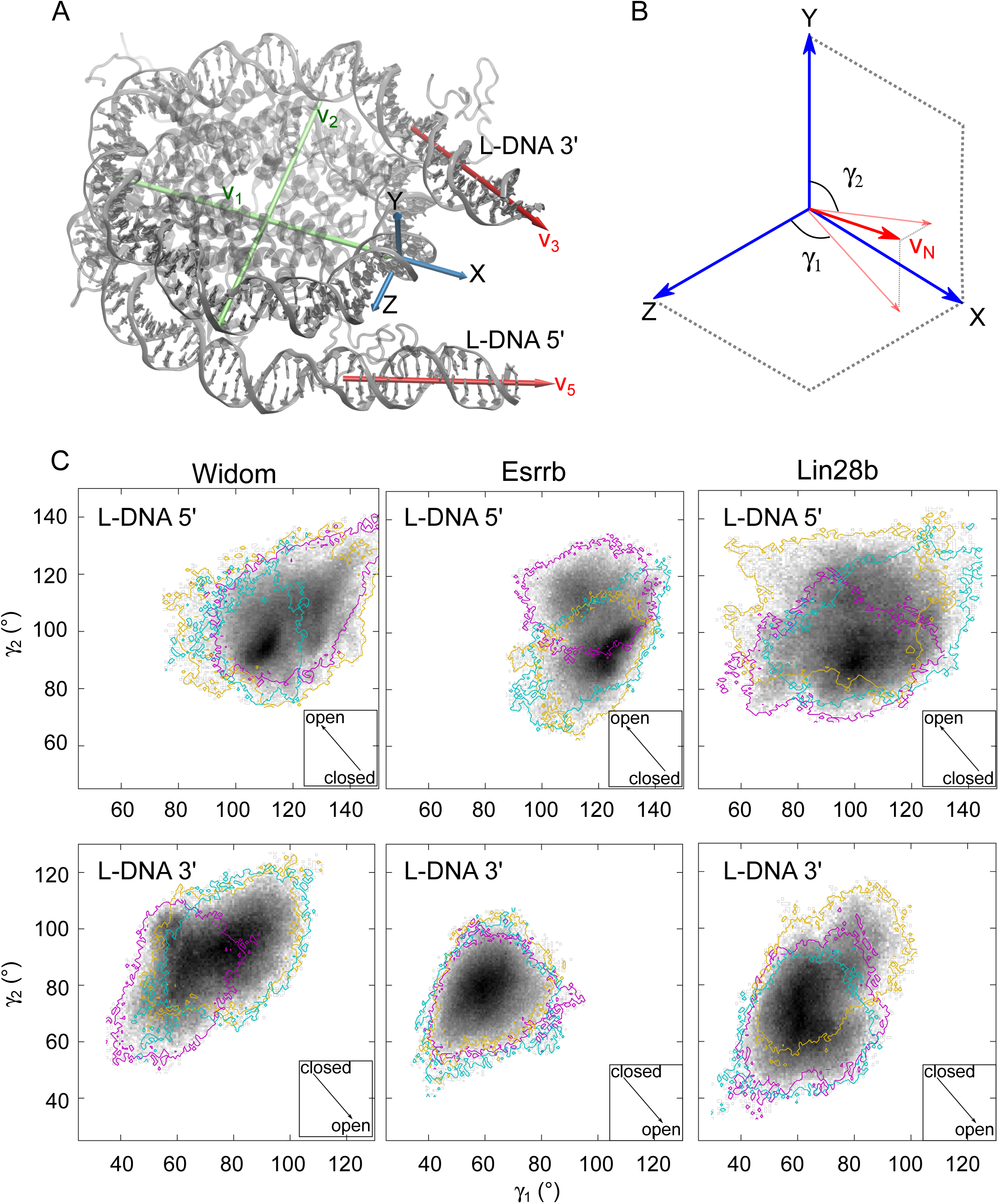
Motions of the linker DNA arms. **(A)** Coordinate system definition: The XYZ reference system was defined as follows (see Methods for details): X was defined along the dyad axis, Y as the cross products between a vector v_1_ defined along the dyad axis and a vector v_2_ defined to be approximately orthogonal to v_1_ and to intersect v_1_ as close as possible to the center of the nucleosome, and Z as the cross product between X and Y. **(B)** Definition of the angles, γ_1_ and γ_2_. γ_1_ is the angle between the projection of the vector v_N_ defined along the helical axis of the linker DNA on the XZ plane, and the Z axis. γ_2_ is the angle formed between the projection of v_N_ on the XY plane and the Y axis. An increase of γ_1_ indicates opening at the 3’ L-DNA, but closing at the 5’ L-DNA, whereas an increase of γ_2_ indicates closing at the 3’ L-DNA and opening it the 5’ L-DNA. **(C)** Two-dimensional histograms showing the sampling of the γ_1 /_ γ_2_ conformational space for both 5’ L-DNA and 3’ L-DNA. The colours (cyan, magenta and gold) indicate the sampling covered by each of the three 1µs simulations. The arrows in the square inserts indicate the direction of the nucleosome opening.

At the 5’ L-DNA of the nucleosome, Widom and Esrrb adopt more closed conformations along γ1 than Lin28b (Fig. 3C, upper row). Although in all three simulations Lin28b sampled more open conformations with lower γ1, in one particular simulation (gold in Fig 3C), this nucleosome adopted wide open conformations characterized by low γ1 and high γ_2_ simultaneously. The largest opening occurred shortly after the start of that simulation and lasted for about 400 ns (Fig. S4A). In general, we observed reversible transitions between closed and open conformations (Fig. S4A). At the 3’ end, the three nucleosomes sampled more similar regions of the conformational space. Lin28b sampled a larger area of the 2D histogram compared to Esrrb but somewhat lower compared to Widom (Fig. 3C, down column). Interestingly, the Widom nucleosome displayed transitions to conformations with high γ1 in two simulations (Fig. S4B). However, during these transitions there was no large opening because the opening along γ_1_ was balanced by closing along γ_2_ (Fig. S4B). Similarly, Lin28b displayed more open conformations along γ_2_ at the 3’ L-DNA arm in the simulation in which the 5’ L-DNA adopted the most closed conformations (third simulation in Fig S4A,B). However, the maximum Pearson correlation coefficient R^2^ between two different angles during a simulation was roughly 0.6, which indicated no correlation between the motions of the two L-DNA arms.

To extract the slow low frequency motions from the simulations, we calculated the essential dynamics using principal component analysis (PCA) and monitored the evolution of the angles γ1 and γ_2_ along the trajectories projected on the first five principal components (PCs). These PCs represent 84.27%, 78.19%, 87.02% of all the motion of the Widom, Esrrb, and Lin28b nucleosomes, respectively. This allowed us to characterize the correlated components of the different types of motions and to separate the uncorrelated motions. PC1 and PC2 described the largest breathing motions in the nucleosomes (Fig. 4). For Widom, both PCs, revealed coupled breathing motions along γ_1_ and γ_2_ at the 5’ end, whereas for both native nucleosomes these motions were uncoupled (Fig. 4 and Fig. S5). Lin28b displayed the largest amplitude breathing at the 5’ end. The Esrrb displayed a larger amplitude breathing than Widom despite its overall lower mobility. At the 3’ end, the motion with the largest amplitude was in Widom along PC1, whereas the breathing of Esrrb had the lowest amplitude and the breathing of Lin28b was restricted along γ_1_. These findings suggest that the pattern of the breathing motions depends on the DNA sequence: in nucleosomes with strong positioning sequences such as the Widom, the breathing along γ_1_ and γ_2_ are coupled and of lower amplitude, while natural nucleosomes have higher uncoupled motions.

**Figure 4.**
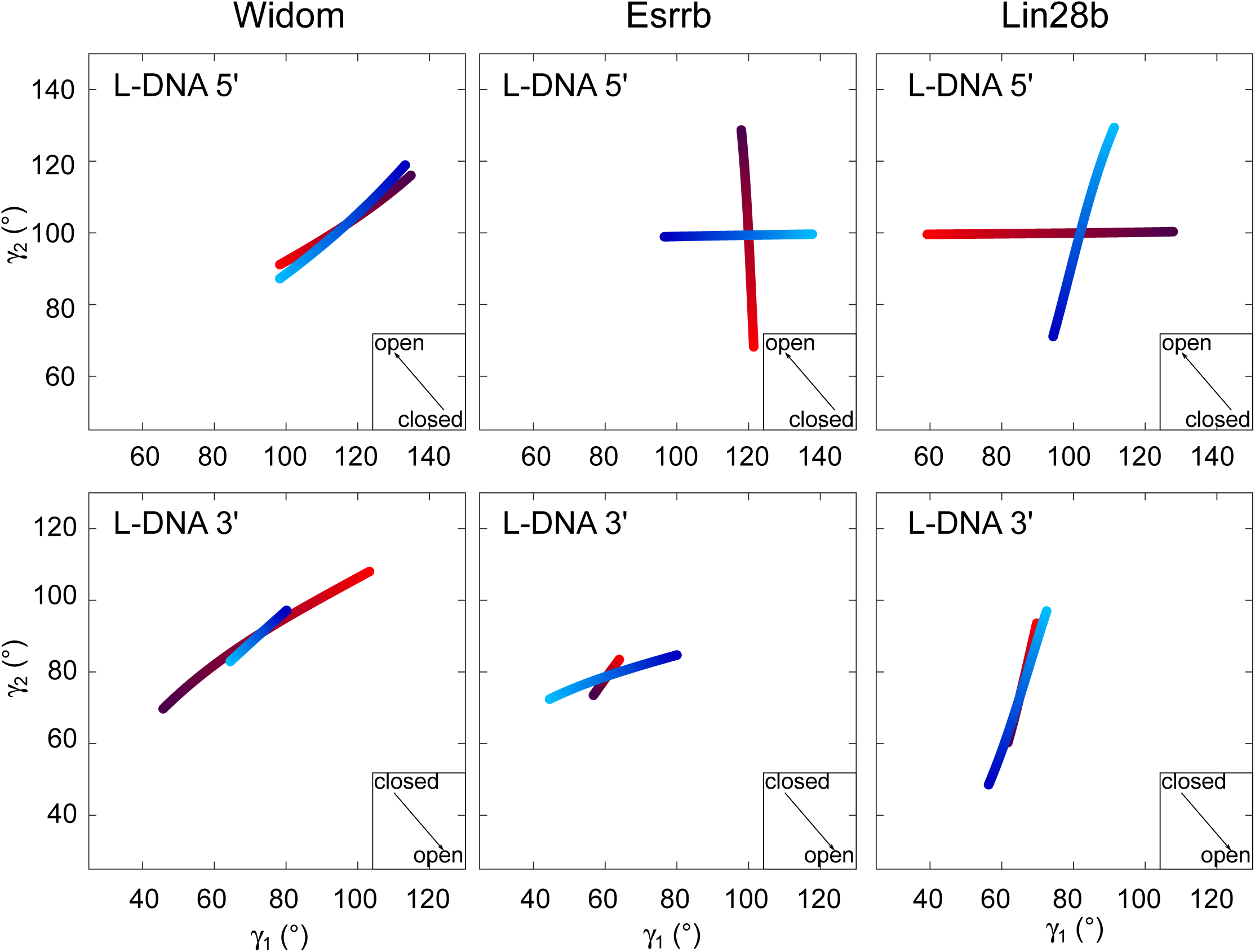
Correlations of nucleosome opening-closing motions. Evolution of the angles γ1 and γ2 in the two first principal components. Following principal component analysis of the ensemble of simulations for each nucleosome, pseudo-trajectories along the two first eigenvectors were generated. Then, the motion of the linker DNA arms was analyzed as described in Figure 3. The motions corresponding to the first and second eigenvectors are in red and blue, respectively. Light and dark colors indicate the low and high amplitudes of the motions, respectively. The arrows in the square inserts indicate the direction of the nucleosome opening.

### Twisting motions in the DNA propagate into the nucleosome core

In addition to breathing, the PCA revealed overtwisting motions of the nucleosomal DNA (Movie S1), which are notoriously difficult to quantify. Previously, these were characterized by monitoring the changes in the contacts of selected arginine residues^23^ or the change of the basepair orientation relative to the nucleosome superhelical axis^15^. Both approaches have important limitations. The first is only suitable for low number of contacts between amino acid side chains and DNA bases. For example, in coarse-grained simulations, often a single contact is established between one amino acid bead and one base pair bead. On the other hand, in atomistic simulations arginines make multiple simultaneous contacts with more than one base pair. The second approach does not provide a clear separation between twisting and other motions. The change in base pair orientation may be due to pure twisting motions or to a base pair rotating during nucleosome breathing. To track the changes in both base pair orientation and position, we developed a new approach by establishing a moving coordinate system for every base pair with the Y axis perpendicular to the helical axis of DNA (Fig 5 A,B, Fig. S6). We then calculated the angle α between Y at a given time and Y at initial time. A concerted, large amplitude motion such as breathing or twisting is characterized by a wide range of values for α. To discriminate between breathing and twisting, we calculated the displacement d of the center of the coordinate system. Breathing motions involve a large d, twisting a moderate d, and no motion, a low d. Therefore, a combination of wide α range and moderate d is a footprint for twisting. We performed the analysis on the pseudo-trajectories along the first five PCs because these display the motion as unidirectional from low to high amplitudes (Fig. S7).

**Figure 5.**
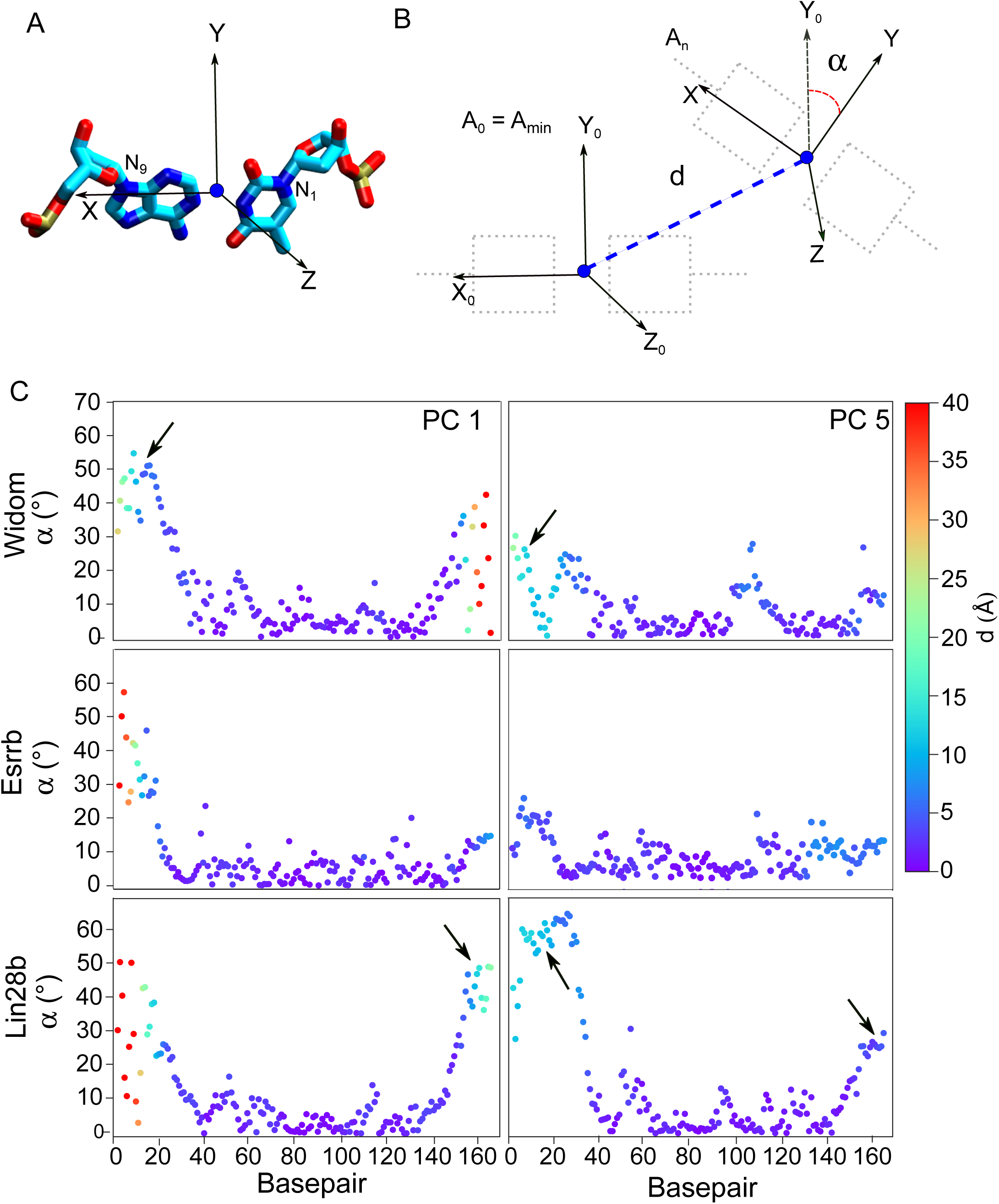
Twisting motions in the nucleosomes. **(A)** Coordinate system definition: The XYZ reference system was defined using the following approach (see also Methods): X is along a vector defined to connect the N1 atom of the pyrimidine with the N9 atom of the purine. Z is along the cross product between X and a vector defined from atom positions to be approximately orthogonal to X. The intersection of this vector with X is the origin of the coordinate system and is near the base pair center. Y is the cross product between X and Z (see also Fig. S6). **(B)** Schematic representation of the motion of a basepair during nucleosome twisting. In each snapshot along the principal component trajectory, the coordinate system is redefined. Then, the displacement d of the current position of the origin from the position at the minimum amplitude and the angle α between the current Y axis and the Y axis at minimum amplitude (Y_0_) were calculated. **(C)** The range of α is plotted for each basepair along the first five principal component trajectories. Each point is colored by the value o*f d.* Twisting motions are characterized by high values of α and low displacement values and are indicated with black arrows.

We focused on the motions along PC1 and PC5 because they described the highest amplitude breathing and twisting. PC5 was the first PC in which twisting was uncoupled from breathing. We observed overtwisting of the DNA in the simulations of Widom and Lin28b (Fig. 5C, Fig. S7) but not in those of Esrrb. For Widom, this motion is revealed mainly by PC1, occurs at the 5’ end, is coupled with breathing and propagates about 30 bp into the core nucleosomal DNA (Fig. 5C, upper left). For Lin28b, the highest amplitude twisting motion was at the 5’ end (Fig. 5C). Along PC5, this motion was uncoupled from breathing and propagated almost 40 base pairs into the core nucleosomal DNA (Fig. 5C lower right, Movie S1). A similar twisting at the 3’ end occurred, but it propagated less (Fig 5C). This large scale twisting originated from a unidirectional overtwisting that occurred in the first simulation of the Lin28b nucleosome, reaching the maximum amplitude after ∼750 ns (Fig. S8). The twisting motions observed in the other simulations (e.g. of Widom) were multi-step bidirectional motions involving partial over- and undertwisting (Fig S8). These findings indicate that the twisting motions also display a higher amplitude in Lin28b compared to Widom and Esrrb.

### Local nucleosome mobility is enhanced around the Oct4 binding sites

All Oct4 binding sites proposed by Soufi et al. (two within Lin28b and one within Esrrb ^5^) (Fig. 6A) are exposed and accessible for Oct4 binding in our nucleosome models despite the low resolution of the MNase data used to build the models (see Methods). In Lin28b, one partial binding site for the POU_HD_ is present and exposed closer to 5’ L-DNA and entire octamer motif with the partial motif recognized being accessible by the POU_S_ is closer to the dyad. From here on, we refer to these sites as the POU_HD_ and POU_S_ binding sites, respectively. For Esrrb, only one partial binding site for the POU_S_ is present and exposed close to the 3’ end.

**Figure 6.**
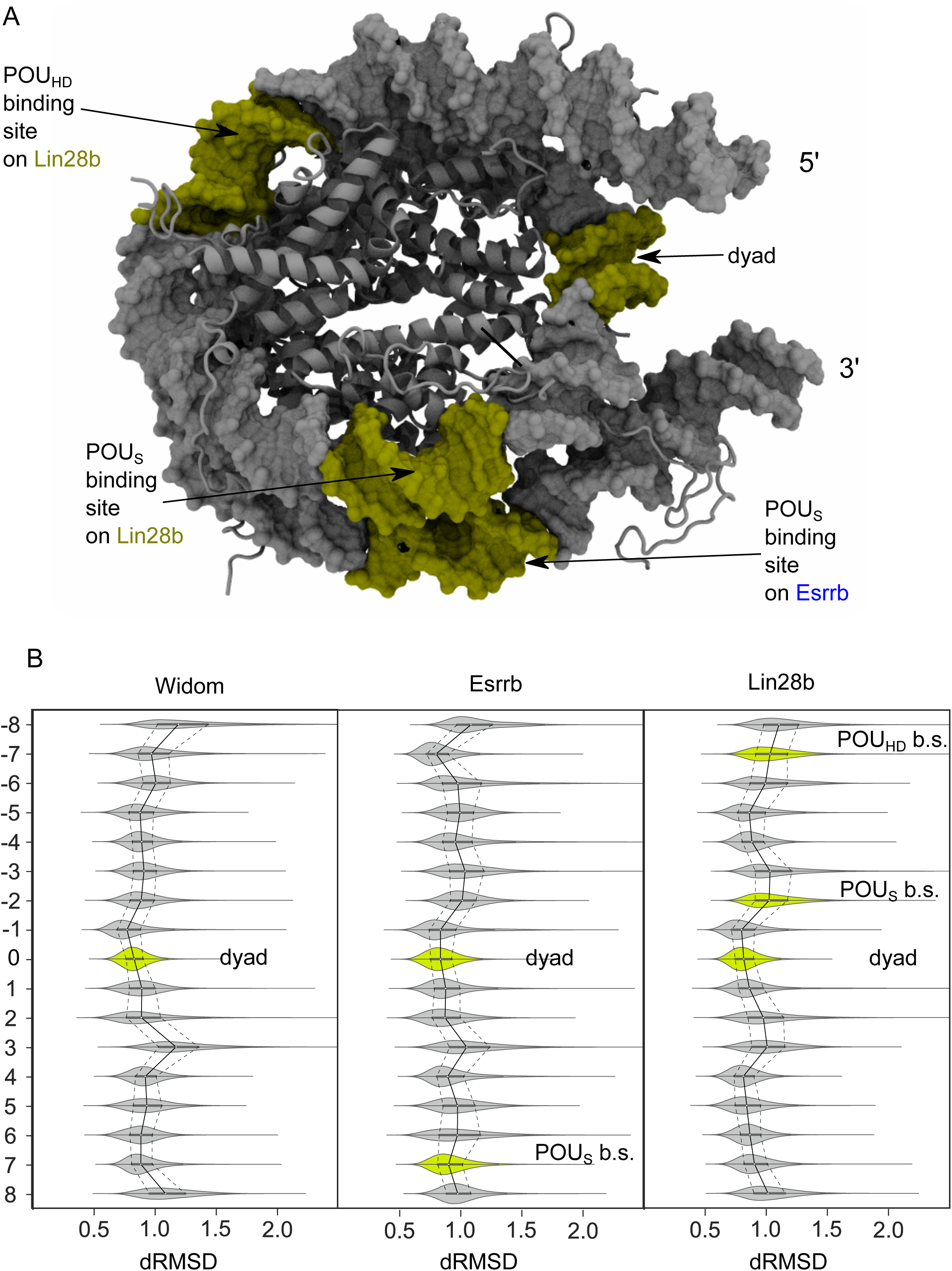
Local flexibility in the nucleosomes. Distributions of dRMSD for different DNA segments. The first segment was the 8 bp region centered on the dyad. The other segments were 8 bp long extending from the dyad segment to the ends of the 146 bp nucleosome core particle. **(A)** Position in the nucleosome of the dyad and the regions containing the Oct4 binding sites (in lime) **(B)** dRMSD distributions for all segments of each nucleosome. The dyad and the Oct4 binding sites are in lime. Lines indicate the median (solid) and the first and third quartiles (dashed).

To investigate whether the Oct4 binding sites are in or close to regions of enhanced local mobility in the nucleosome (Fig 1), we divided the DNA in 8 base pair fragments starting from the dyad. For each fragment, we calculated the dRMSD using the average structure of the fragment DNA backbone from all simulations as reference. We visualized the results as violin plots that revealed both the statistical median and quartiles, and the histograms of the distribution of dRMSDs. As expected, the dyad fragment was the region with the lowest mobility (Fig. 6B). Interestingly, in both Esrrb and Lin28b, both regions next to the dyad showed increased local mobility, whereas in Widom, only the region at the 3’ side of the dyad was more mobile. In addition, the oscillatory pattern of the local mobility in which more mobile regions alternate with less mobile regions was more pronounced in the native nucleosomes (Fig 6B). Both in Lin28b and Esrrb, the Oct4 binding sites previously identified experimentally^5^ were located either in or next to regions of increased local mobility, suggesting that the accessibility to these binding sites is facilitated by increased local nucleosome mobility.

### Alternative configurations of Oct4 form stable complexes with nucleosomes

Finally, to reveal the structural basis for Oct4-nucleosome recognition we modeled the structure of Oct4 bound to Lin28b using the experimental data available from Soufi et al^5^. The two binding sites for Oct4 on Lin28b were proposed to be recognized by the POU _S_ and POU_HD_, respectively, from biochemical, footprinting, and ChIP-Seq experiments.

We confirmed that the recognition of the entire octamer motif by Oct4 with both domains bound in the configuration known from the crystal structure of Oct4 on free DNA (Fig 7A, left structure) was incompatible with nucleosome binding on all proposed binding sites due to steric clashes between one domain and the histones (Fig 7B-D left structures). Therefore, we applied alternative Oct4 configurations. First, we used a configuration found in the structure of an Oct4 homodimer bound to the palindromic MORE motif (MORE configuration)^10^. In this configuration, both the POU_S_ and POU_HD_ are bound on the same side of the DNA (Fig 7A, middle structure). Second, we used different Oct4 configurations generated from MD simulations of free Oct4 (Fig 7A, right structure) (MD configurations). We simulated the sequence specific binding of one Oct4 domain in multiple step restrained MD equilibrations by applying restraints to enforce the formation of the interactions with DNA bases known from the Oct4-DNA structure (PDB ID 3L1P). In the last step of the equilibrations, we released these restraints. Then, we monitored the stability of the Oct4- DNA interactions in unrestrained, 100 ns long MD simulations started from those equilibrations in which the interactions were maintained (Table 2) (see Methods for details). We evaluated the stability of the Oct4-nucleosome configurations with the following criteria: the number of stable contacts with the DNA, and the presence of specific hydrogen bonds between specific amino acids and DNA bases (Q44-A in the POU_S_ and N143-A in the POU_HD_^6^,^31^). We expected the protein-DNA interactions to break quickly at the beginning of the simulations for any invalid Oct4 binding mode.

**Table 2:** Oct4-DNA contacts in Oct4-nucleosome complexes.

| Binding Site | Oct4 configuration | Simulation step | # stable contacts<sup>†</sup><br>POU<sub>S</sub> – DNA bases | # stable contacts<sup>†</sup><br>POU<sub>S</sub> – DNA backbone | # stable contacts<sup>†</sup><br>POU<sub>HD</sub> – DNA bases | # stable contacts<sup>†</sup><br>POU<sub>HD</sub> – DNA backbone | % Time with selected contact<sup>‡</sup> present |
| --- | --- | --- | --- | --- | --- | --- | --- |
| POU<sub>S</sub> | MD | Eq01* | 67 | 38 | 0 | 0 | 92,40% |
|  |  | Eq02 | 14 | 43 | 0 | 0 | 0% |
|  |  | Eq03 | 30 | 51 | 0 | 0 | 0,52% |
|  |  | Prod | 61 | 50 | 0 | 0 | 95,32% |
|  | MORE | Eq01 | 69 | 47 | 4 | 23 | 85,80% |
|  |  | Eq02 | 47 | 71 | 0 | 6 | 6,56% |
|  |  | Eq03* | 69 | 77 | 0 | 1 | 99,64% |
|  |  | Prod | 65 | 54 | 2 | 7 | 98,73% |
| POU<sub>HD</sub> 1<br>(3'-5') | MD1 | Eq01 | 0 | 0 | 31 | 8 | 0% |
|  |  | Eq02 | 0 | 2 | 3 | 13 | 0% |
|  |  | Eq03 | 0 | 0 | 21 | 15 | 0% |
|  |  | Prod | - | - | - | - | - |
|  | MD2 | Eq01* | 1 | 5 | 41 | 44 | 65,52% |
|  |  | Eq02 | 1 | 5 | 17 | 20 | 0% |
|  |  | Eq03 | 0 | 0 | 12 | 26 | 0% |
|  |  | Prod | 0 | 0 | 5 | 11 | 19,84% |
| POU<sub>HD</sub> 2<br>(5'-3') | MD | Eq01 | 0 | 0 | 46 | 47 | 69,80% |
|  |  | Eq02 | 0 | 0 | 34 | 37 | 87,08% |
|  |  | Eq03* | 0 | 0 | 42 | 16 | 88,16% |
|  |  | Prod | 0 | 0 | 34 | 17 | 63,98% |
|  | MORE | Eq01* | 18 | 47 | 48 | 37 | 97,68% |
|  |  | Eq02 | 23 | 51 | 50 | 50 | 88,68% |
|  |  | Eq03 | 16 | 50 | 42 | 51 | 95,84% |
|  |  | Prod | 2 | 35 | 31 | 27 | 85,43% |

Abbreviations: *Eq* = Equilibration (10,25 ns); *Prod* = Production (100 ns)
\* Equilibration chosen for production
† Stable contact: Atoms at a distance < 4.5 Å for at least 75% of the simulation
‡ Selected contacts: Q44-A for the POU<sub>S</sub> and N143-A for the POU<sub>HD</sub>. Contact was considered present if donor-acceptor distance was smaller than 4 Å

Binding in the POU_S_ binding site was stable in 2 of the 3 equilibrations with the Oct4 MORE configuration and in 1 of the 3 equilibrations with the MD configuration. The interactions remained stable during the 100 ns simulations (Table 2, Figure 7B, middle and right structures, Movie S2). In these complexes, the POU_HD_ hovered unspecifically on the same gyre of the nucleosome. For the POU_HD_ to reach the second gyre, a major conformational change of the linker between the 2 domains must occur. Interestingly, some models suggest that the 2 domains may recognize their sequence specific binding sites simultaneously if the linker was in an extended conformation (Figure 7B, right structure).

**Figure 7.**
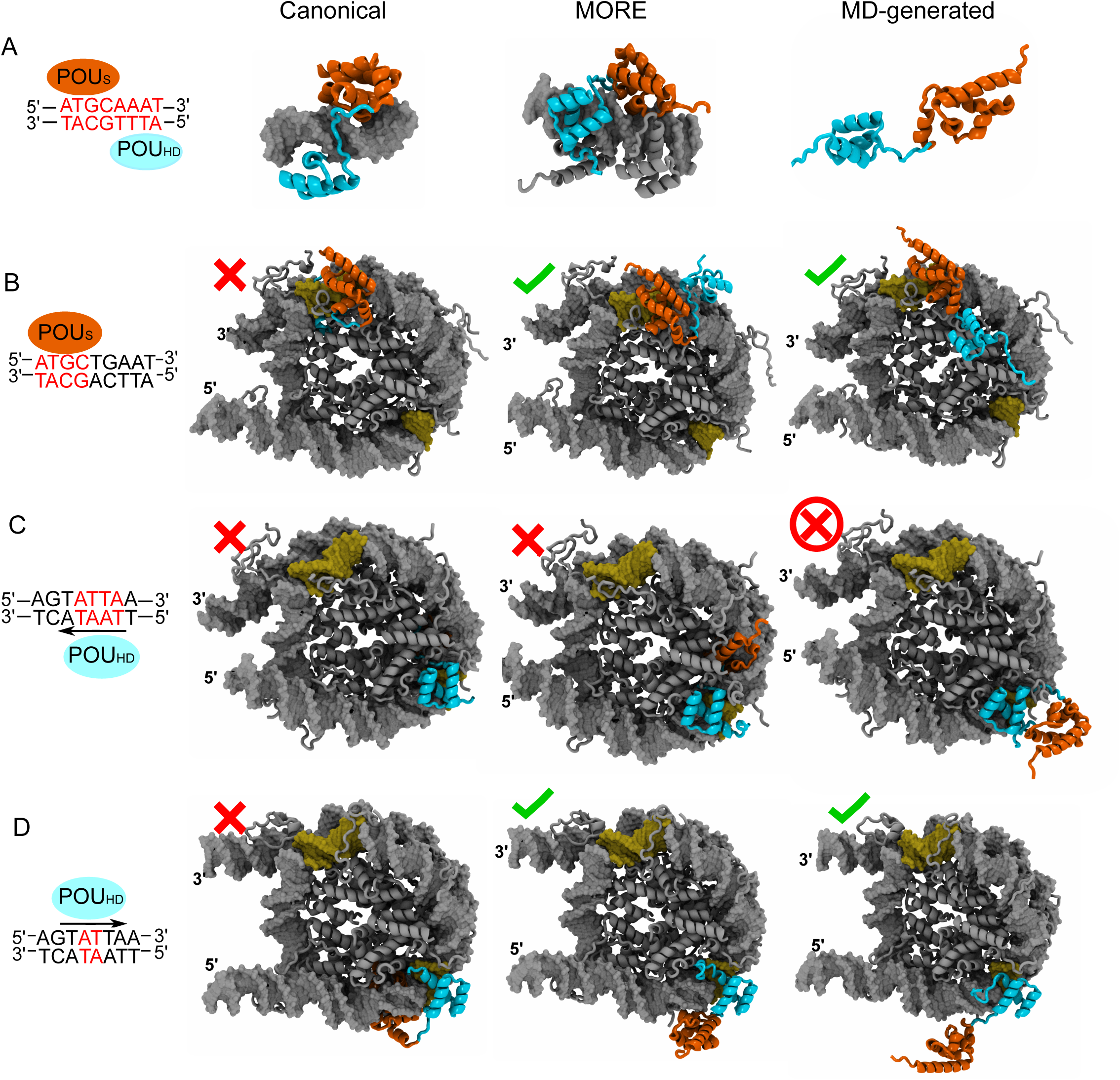
Structural basis for Oct4-nucleosome recognition. **(A)** Structures of Oct4 used to build Oct4-nucleosome complexes. From left to right a schematic representation of the canonical configuration found in the crystal structure of an Oct4 dimer bound to palindromic DNA^6^ is followed by structural views of the canonical configuration, MORE homodimer configuration (originally found in the crystal structure of Oct1-Oct1-DNA complex, and modelled in our previous work^10^), and an example of a configuration obtained from MD simulations of free Oct4. The recognized bases are highlighted in red. **(B,C,D)** Oct4-nucleosome complexes. From left to right a schematic representation indicating which domain is bound in a sequence-specific manner is followed by structural models of Oct4-nucleosome complexes built using the canonical, MORE, and MD-generated configurations. **(B)** POU_S_ is bound sequence specifically. **(C**, **D)** POU_HD_ is bound sequence-specifically either in an orientation as observed in structures of Oct4-DNA complexes **(C)** or in an orientation in which an AT base pair step on the opposite strand is recognized **(D)**. The black arrows indicate the orientation of the POU_HD_ binding. The POU_S_ and POU_HD_ are in orange and cyan cartoons, whereas the nucleosome core is in grey cartoons.

Binding in the POU_HD_ binding site may occur in 2 different orientations. For a complete recognition of the typical “TAAT” site, the POU_HD_ has to be bound in 3’-5’ orientation (Figure 7C, left drawing). The MORE configuration was not compatible with such a binding mode because of steric clashes between the POU_S_ and the histones (Fig 7C, middle structure). Moreover, in all simulations started with MD configurations, the interactions of the POU_HD_ residues with DNA bases were not stable (Table 2, Figure 7C right structure, Movie S3). Therefore, we concluded that this binding mode is not valid. In an alternative binding mode, the POU_HD_ recognizes sequence-specifically only an “AT” dinucleotide on the 5’-3’ strand (Fig. 7D, left drawing). Interestingly, both the MORE and MD configurations of Oct4 were compatible with this binding mode forming stable complexes with typical POU_HD_-DNA interactions maintained during the simulations (Table 2, Figure 7D middle and right structures, Movie S4). This suggests that the POU_HD_ is able to bind to the nucleosomal DNA in different orientations comparing to free DNA. In these models, the POU_S_ remained unspecifically close to DNA on the same gyre as the POU_HD_. Again, the binding of the 2 domains to different gyres may occur only upon a large conformational transition in the linker region.

These findings show that alternative but not all configurations of Oct4 with either the POU _S_ or the POU_HD_ bound to sequence specific sites formed stable interactions with the nucleosomes on the simulated time scale. This demonstrates that Oct4 binding is compatible with the nucleosome curvature and the histone-DNA interactions and that MD simulations are powerful in identifying configurations and orientations of multidomain TFs bound to nucleosomes.

## Discussion

Here we revealed the nucleosome dynamics that mediate the binding of the pioneer transcription factor Oct4 to DNA wrapped in nucleosomes and probed for the structural basis of Oct4-nucleosome recognition using microsecond time scale MD simulations. We first studied the dynamics of 3 nucleosomes, 2 with natural DNA sequences proposed to be bound by Oct4 during cellular reprogramming to pluripotency^4^ and 1 with a strong positioning sequence and no TF binding site. Whereas the dynamics of artificial nucleosomes has been previously studied with atomistic simulations^14–16^, the sequence dependence of these dynamics has not been explored. We found significant differences between the dynamics of the 3 nucleosomes, the nucleosome with most TF binding sites showing the largest amplitude of breathing and twisting motions. This is in agreement with previous studies that revealed the more dynamic nature of natural nucleosomes^13,22,28^. Based on our findings, we propose that nucleosomes with multiple TF binding sites are more mobile and less compact. However, we also found that one of the natural nucleosomes we studied was at least as rigid as the artificial nucleosome, suggesting that the mobility of natural nucleosomes varies significantly.

Our findings were in agreement with *in vitro* experiments in which temperature-induced nucleosome disassembly was performed. The most mobile nucleosome in the simulations also disassembled at the lowest temperature. Moreover, the natural nucleosome that was more rigid in the simulations had different disassembly profiles in the experiments (Fig S1). In one experiment, the disassembly temperature of this nucleosome was highest, in agreement with the rigidity observed in the simulations. This suggests that natural nucleosomes may exist in different states, some more dynamic than others. In such cases, the simulations are unlikely to describe the transition between different states due to insufficient sampling and large energy barriers involved.

Nucleosome motions are particularly difficult to study in atomistic simulations due to the long time scales involved^32,33^. Despite the lack of full sampling convergence, we observed a good overlap between independent simulations for nucleosome breathing (Fig 3) and we achieved a wider conformational sampling by combining independent simulations of the same species into ensembles. This enabled us to study both breathing and twisting motions in the nucleosomes and to conclude about the sequence dependence of these motions at a level of detail not feasible in coarse grained simulations, which were previously used to study nucleosome dependent dynamics^22,28^. The atomic resolution provides the means to study the determinant modulators of nucleosome dynamics such the motion of histone tails or the role of anchoring histone residues^15^. Ultimately, a combined technique that would allow on-the-fly conversion between different representations during the simulation may be the most powerful in revealing dynamics of large structures such as the nucleosome or larger chromatin fibers.

Our findings suggest that the nucleosomes with multiple TF binding sites are more dynamic and that TF binding sites are located in or near regions of increased local mobility in these nucleosomes. This suggests that binding sites located in dynamic nucleosomes are more prone to be recognized and bound by TFs. Therefore, the global and local mobility of the DNA wrapped around the core histones may function as mediator for binding of proteins to nucleosomes. We have previously shown that nucleosome dynamics play a role in the binding of the linker histone to nucleosomes^18^. Based on these findings, we propose that alongside DNA sequence and post-translational modifications of histone tails, nucleosome dynamics provide an additional platform for protein-nucleosome recognition.

Ultimately, we provide here a structural basis for Oct4-nucleosome recognition. This is particularly important because to date no experimental structure of a TF bound to nucleosome has been resolved. Understanding this interaction at atomic resolution is to provide novel approaches to modify the functional outcome of TF binding to DNA. For example, enhancing nucleosome binding of selected pioneer factors may lead to faster and more efficient chromatin decompaction, which in turn may have profound effects on gene regulation programs. In the case of Oct4, we first confirmed the partial binding site recognition model proposed by Soufi et al^5^. Then, we revealed alternative configurations of Oct4-nucleosome complexes that maintained the Oct4-DNA interactions during 100 ns long MD simulations. Our findings suggest that the nucleosome curvature and DNA-histone interactions do not interfere with TF binding. Because all our simulations were started with intact nucleosomes, we cannot exclude a mechanism for TF-nucleosome binding that involves an initial partial decompaction and unwrapping of the nucleosome followed by recognition of the binding sites as in free DNA.

Furthermore, we showed that MD simulations are powerful to identify configurations that are incompatible with nucleosome binding. Longer classical MD simulations of TF-nucleosome complexes and comparing with similar-length simulations of free nucleosomes will provide a framework for studying the role of TF binding to nucleosome dynamics. For this, enhanced sampling techniques may be required to capture larger scale motions and longer time scales^34^.

## Methods

### Nucleosome Modeling

The 147 bp Widom *Drosophila melanogaster* nucleosome was built by homology modeling using as templates the structures of the *D. melanogaster* nucleosome core (Protein Data Bank [PDB] ID 2PYO), the structure of the Widom 601 Nucleosome Core Particle (PDB ID 3LZ0) and the high resolution structure of the nucleosome core particle including tails (PDB ID 1KX5). Using Modeller (https://salilab.org/modeller/), 100 homology models were generated using a “slow” optimization protocol followed by a “slow” molecular dynamics based refinement protocol. The model with the lowest Discrete Optimized Protein Energy was then selected. Then fragments of B-DNA generated with NAB (available in Ambertools 18^35^) were added to each end to generate the 168 bp nucleosome.

The Esrrb and Lin28b enhancer sequences with different Oct4 binding profiles were selected from data by Soufi et al.^4^. The Lin28b sequence, was selected as a representative of a nucleosome with multiple TF binding sites, whereas the Esrrb sequence was chosen for having a single binding site of Oct4. To model these natural nucleosomes, we used the MNase data that was used by Soufi et al. to superimpose on Oct4 binding data obtained from ChIPSeq^36^. We substituted the Widom sequence with the Lin28b and the Esrrb sequences using the “swapna” function in Chimera (https://www.cgl.ucsf.edu/chimera/) upon fixing the center of the MNase peak to be the dyad point. Since the MNase data did not have single bp resolution, the Lin28b sequence was shifted by one base pair to obtain a conformation with a better exposure of the Oct4 binding sites.

### Modeling of the Oct4-nucleosome complex

We used the following structures to model Oct4 bound to nucleosomes: (i) Oct4 bound in the canonical configuration as found in the structure of the homodimer bound to the palindromic PORE motif^6^, (ii) Oct4 bound as homodimer to the partially palindromic MORE motif, (iii) free Oct4 from MD simulations. The canonical configuration of Oct4 was obtained by modeling the human Oct4 using as a template the final snapshot from the previously published trajectories of Oct4 bound to free DNA^9^. The MORE configuration was obtained by stripping one monomer from the first snapshot after equilibration of the published trajectories of Oct4 bound as a homodimer in the MORE configuration^10^. Finally, the MD configurations of Oct4 were obtained after 100 ns of MD simulations of Oct4, starting from the canonical configuration upon stripping the DNA (see details of the simulation in molecular dynamics simulations subchapter).

To model the binding of Oct4 to the nucleosome, we assumed that the interactions between Oct4 residues and DNA bases (direct readout) will be the same as in the free DNA. This assumption was based on ChIP-Seq data that revealed the same recognition motif for Oct4 on free and wrapped DNA. First, we superimposed the Oct4 binding sites from the free and nucleosomal DNA, using only bases that form direct readout interactions with Oct4. Then, we removed the free DNA from the resulting model. For the MD configurations, we used the hypothetical Oct4-nucleosome model with Oct4 in the canonical configuration as template on which we superimposed the selected MD configuration. Snapshots with no clashes between Oct4 and the histones or the DNA were selected for further MD simulations.

### Molecular dynamics simulations

The procedure below was applied for the simulations of free Oct4 alone, free nucleosomes, and Oct4-nucleosome complexes. Each species were solvated in a truncated octahedron box of TIP3P water molecules, with a layer of at least 12 Å of water around the solute in any direction. Na^+^ ions were added to counter the negative charges. KCl was added to the system up to a concentration of 150 mM. The solvated systems were then optimized by energy minimization with the AMBER software^35^. After that, the systems were equilibrated in NAMD^37^ for 10.25 ns using restraints that were removed in a stepwise manner, using a protocol described in detail in our previous simulations of protein-DNA systems^10^. After equilibration, the system were simulated using NAMD in the isobaric-isothermic (NPT, p = 1 atm, T = 300 K) ensemble, using a standard protocol with Langevin dynamics to control the temperature and Nose Hoover – Langevin Piston to control pressure. The Merz ion parametrization, the ff14SB^38^ and the parmbsc1 forcefields^39^ were used for ions, protein and DNA respectively. Oct4 alone was simulated 5×100 ns to obtain conformations compatible with nucleosome binding. Each nucleosome was simulated for 3×1µs, whereas the Oct4-nucleosome structures were submitted to three independent equilibrations and one equilibration for every configuration was simulated for 100 ns.

### Analysis of nucleosome dynamics

The MD trajectories were processed by imaging and performing an RMS-fit of the heavy atoms of the histone core (excluding histone tails) to the minimized structure of the Widom nucleosome. Atomic fluctuations of the heavy atoms of the DNA were calculated and averaged by residue using cpptraj^40^. The fluctuations were squared and weight by 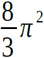 to obtain a value similar to a crystallographic B-factor. To facilitate the comparison, the average value of mobility of the DNA core (146 bp) was calculated, and all B-factors were normalized by that factor. Then, DNA residues were colored using VMD^41^, with blue and red indicating normalized B-factors smaller than 1 or bigger than 1, respectively.

Distance Root Mean Square Deviations (dRMSD), the RMSD matrices and the radius of gyration were also calculated using cpptraj. In the fitted trajectories, the dRMSD and the mass-weighted radius of gyration was obtained for the heavy atoms of the DNA. Calculations were done for DNA fragments of 168 bp, 146 bp, 138 bp, and 120 bp. To obtain the fragments smaller than 168 base pairs, we excluded 11, 15, and 24 base pairs from each L-DNA from the analysis. The average structure of the DNA backbone of all three 3 µs-ensembles was used as reference to prevent bias coming from the reference structure. Similarly, the RMSD matrices were obtained by calculating the RMSD of the heavy atoms of the DNA between each 2 snapshots for the different DNA fragments. To characterize the regions of local mobility, the DNA was fragmented in regions of 8 bp each, starting from the dyad, and the dRMSD was calculated for each fragment, using again the average of the DNA backbone of all three ensembles as reference.

To characterize the breathing motions of the nucleosomes, we applied the procedure described by Öztürk et al.^18^. First, we defined a coordinate system XYZ with the origin on the dyad. X was defined along the dyad axis, Y as the cross products between a vector v_1_ defined along the dyad axis and a vector v_2_ defined to be approximately perpendicular to v_1_ and to intersect v_1_ as close as possible to the center of the nucleosome, and Z as the cross product between X and Y. Then, the angles γ_1_ and γ_2_ were defined as the angle between the projection of the vector v_N_ defined along the helical axis of the linker DNA on the XZ plane, and the Z axis, and γ_2_ as the angle formed between the projection of v_N_ on the XY plane and the Y axis.

Principal Component Analysis was performed in cpptraj using the non-hydrogen atoms of the DNA and the rigid part of the histones. The covariance matrix was calculated and diagonalized to extract the first 25 eigenvectors and eigenvalues. Then, the trajectory was projected on the first 5 modes and the minimum and maximum projection values for each mode were extracted. Finally, pseudo-trajectories along each mode were generated in order to analyze the motions of the system.

To quantify the twisting motions of the nucleosomes, and distinguish them from other motions, a basepair-centered method was developed (Figure S6). For every basepair, a vector X was defined along the line that connects the N1 atom of the pyrimidine with the N9 of the purine. Then, a vector connecting the geometrical center of the C_2_ atoms of both bases and the geometrical center of the C_4_ atoms of both bases was defined. Z was set along the cross product between this vector and X. The origin O of the coordinate system was defined at the intersection between this vector and X which is near the basepair center. The coordinate system was updated in each snapshot, the updated origin being O’ and the updated axes X’, Y’, and Z’. Finally, we calculated the angle formed between Y and Y’ for each base pair and the displacement d as the distance between O and O’.

### Analysis of Oct4-nucleosome complexes

To characterize the stability of the Oct4-nucleosome complexes, we used the last, unrestrained step of the equilibrations (5 ns) and the production runs (100 ns). The binding of Oct4 was evaluated using two criteria. On one hand, the number of stable contacts between Oct4 and the DNA (defined as the number of Oct4 non-hydrogen atoms closer than 4.5 Å to the DNA that are maintained for at least 75% of the simulated time) was determined using cpptraj. As direct and indirect readout interactions, we considered the contacts with the DNA bases and backbone, respectively. On the other hand, we monitored the specific hydrogen bonds formed by Q44 of the POU_S_ and N143 of POU_HD_ with adenine bases. These are direct readout interactions selected based on the Oct4- DNA structure^6,31^. The % of simulation time in which the donor – acceptor distance was smaller than 4 Å was calculated using VMD.

### Mononucleosome preparation

Full-length, *Drosophila melanogaster* histones were individually purified and refolded in a ratio of 1:1:1:1 according to the protocol published by Luger et al.^42^. Unlabeled, high- concentration nucleosomes were assembled using the salt gradient dialysis method described in the same publication. Assembled nucleosomes were purified from free DNA and free histones using a 10-30% sucrose gradient spun at 30 k x g and 4°C in a Beckman Coulter Optima L-100 XP swing bucket rotor (SW-41) for 18 hours. Fractions were collected from the bottom of the gradient, screened on native gels, pooled, and quantitated by densitometry using a DNA standard curve and Quantity One® software (Bio-Rad).

### Thermal stability assay

Nucleosome thermal stability was assessed according to the protocol outlined in Taguchi et al.^30^. Briefly, nucleosomes were diluted to 2.5 µM in low salt buffer (25 mM Hepes pH 7.8, 50 mM NaCl, 0.5 mM EDTA, and 10% glycerol) along with SYPRO® Orange diluted 1:1000 (Sigma Cat. # S5692) in a 20 µL final reaction volume. Reaction components were carefully mixed, briefly spun down, and immediately measured. Samples were analyzed in triplicate over the temperature range 25-95°C in a 96-well microplate using the TAMRA detection settings on an ABI 7900HT. Wells containing water or 2.5 µM BSA standard (Bio-Rad Cat. # 5000206) were included as controls. Raw data were collected from the channel with the maximum signal among all the samples, normalized, and replotted to find the T_max_.

## Acknowledgments

We thank Abdenour Soufi for providing the nucleosome sequences and for discussions. We’d also like to thank Peter Becker and Catherine Regnard at the LMU in Münich for nucleosome assembly training and reagents. This project was supported by funds of the Max Planck Society. For high performance computing resources, JH and VC thank the Max Planck Society and the Gauss Centre for Supercomputing e.V. (www.gauss-centre.eu; project ID 12622, STRUCNUCREC running on the GCS Supercomputer SuperMUC at Leibniz Supercomputing Centre (www.lrz.de). JH is part of the International Max Planck Research School-Molecular Biomedicine, Münster, Germany.

## Author contributions

JH designed, performed, and analyzed the simulations and wrote the manuscript; CM designed, performed, analyzed the experiments and contributed discussions and writing; HRS contributed to study design, discussion, and writing; VC designed the simulations, contributed analysis and discussion, and wrote the manuscript.

---

**Supplementary Figure 1. Spyro Orange Thermal Stability Assay (see also Table 1).** Fluorescence readings of the 3 replicas of the Spyro Orange Thermal Stability Assay. For each sequence, the fluorescence is normalized, and the temperature at which there is a peak of fluorescence is marked with a vertical dotted line as the nucleosome dissociation temperature.

**Supplementary Figure 2. RMSD matrices for different DNA regions (see also Figure 1).** From the 3 µs ensemble of simulations for every nucleosome, snapshots were saved every 5 ns. After fitting the snapshots to the heavy atoms of the structured part of the histones (excluding the histone tails), the RMSD of the heavy atoms of the DNA between each 2 snaphots was calculated for the following DNA regions: **(A)** The entire simulated DNA (168 bp). **(B)** The core nucleosomal DNA (146 bp) obtained by excluding 11 bp of linker DNA at each end. **(C)** Nucleosomal DNA obtained by excluding 15 bp of DNA at each end (138 bp). **(D)** Nucleosomal DNA corresponding to the complete inner DNA turn and half of the outer DNA turn obtained by excluding 23 bp of DNA at each end (120 bp). The individual simulations are separated by black lines.

**Supplementary Figure 3. Nucleosome flexibility and compaction (also see Figure 2).** The evolution of the dRMSD (top) and RoG (bottom) of the DNA over time, for the three simulations of the three nulceosomes, is shown for the following DNA regions: **(A)** The entire simulated DNA (168 bp). **(B)** The core nucleosomal DNA (146 bp) obtained by excluding 11 bp of linker DNA at each end. **(C)** Nucleosomal DNA obtained by excluding 15 bp of DNA at each end (138 bp). **(D)** Nucleosomal DNA corresponding to the complete inner DNA turn and half of the outer DNA turn obtained by excluding 23 bp of DNA at each end (120 bp). The individual simulations are separated by vertical black lines.

**Supplementary Figure 4. Time evolution of nucleosome opening-closing motions (also see Figure 3).** Time series for the γ angles described in Figure 3. Next to every plot, a histogram indicating the total sampling of the angle values during the simulations is shown **(A)** γ_1_ of LDNA-5’ **(B)** γ_2_ of LDNA-5’ **(C)** γ_1_ of LDNA-3’ **(D)** γ_2_ of LDNA-3’. The individual simulations are separated by vertical black lines.

**Supplementary Figure 5. Opening-closing motions along the first five principal components (also see Figure 4).** The motion of the linker DNA arms was analyzed using the coordinate system established in Figure 3 on the pseudo-trajectories generated along modes 1 to 5. A lighter tone indicates the lowest amplitude of the mode, with darker tones corresponding to the higher amplitudes. The arrows in the square inserts indicate the direction of the nucleosome opening

**Supplementary Figure 6 (also see Figure 5). Definition of the basepair-centered coordinate system.** This was used to characterize the twisting motions shown in Figure 5 and Supplementary Figure 7. X was defined along the vector connecting the N_1_ atom of the pyrimidine with the N_9_ atom of the purine. Then, a vector connecting the geometrical center of the C_2_ atoms of both bases and the geometrical center of the C_4_ atoms of both bases was defined. Z was set along the cross product between this vector and X. The origin of the coordinate system is at the intersection between this vector and X which is near the basepair center. Y was set as the cross product between X and Z.

**Supplementary Figure 7 (also see Figure 5). Twisting motions along the first 5 principal components.** The plots show the range the angle *α*, colored by the mean displacement *d*, calculated as defined in Figure 5 on the pseudo-trajectories generated along the first five eigenvectors of the PCA analysis for each nucleosome. The occurrence of twisting motions is indicated by black arrows.

**Supplementary Figure 8 (also see Figure 5). Projection of trajectories on the fifth eigenvector of the ensemble of simulations of the Li2n8b nucleosome.** The individual simulations are separated by vertical black lines.

**Supplementary Movie S1. Pseudo-trajectory of Lin28b PC5 reveals unidirectional overtwisting of the nucleosomal DNA**

**Supplementary Movie S2. Stable sequence specific interactions between Oct4 and the Lin28b nucleosome in the POU**_**S**_ **binding site.**

**Supplementary Movie S3. Unstable sequence specific interactions between Oct4 and the Lin28b nucleosome in the POU**_**HD**_ **binding site when the POU**_**HD**_ **is bound in the 3’-5’ orientation.**

**Supplementary Movie S4. Stable sequence specific interactions between Oct4 and the Lin28b nucleosome in the POU**_**HD**_ **binding site when the POU**_**HD**_ **is bound in the 5’- 3’ orientation.**

