## Supplementary Figures for "Nucleosomal DNA dynamics mediate Oct4 pioneer factor binding"

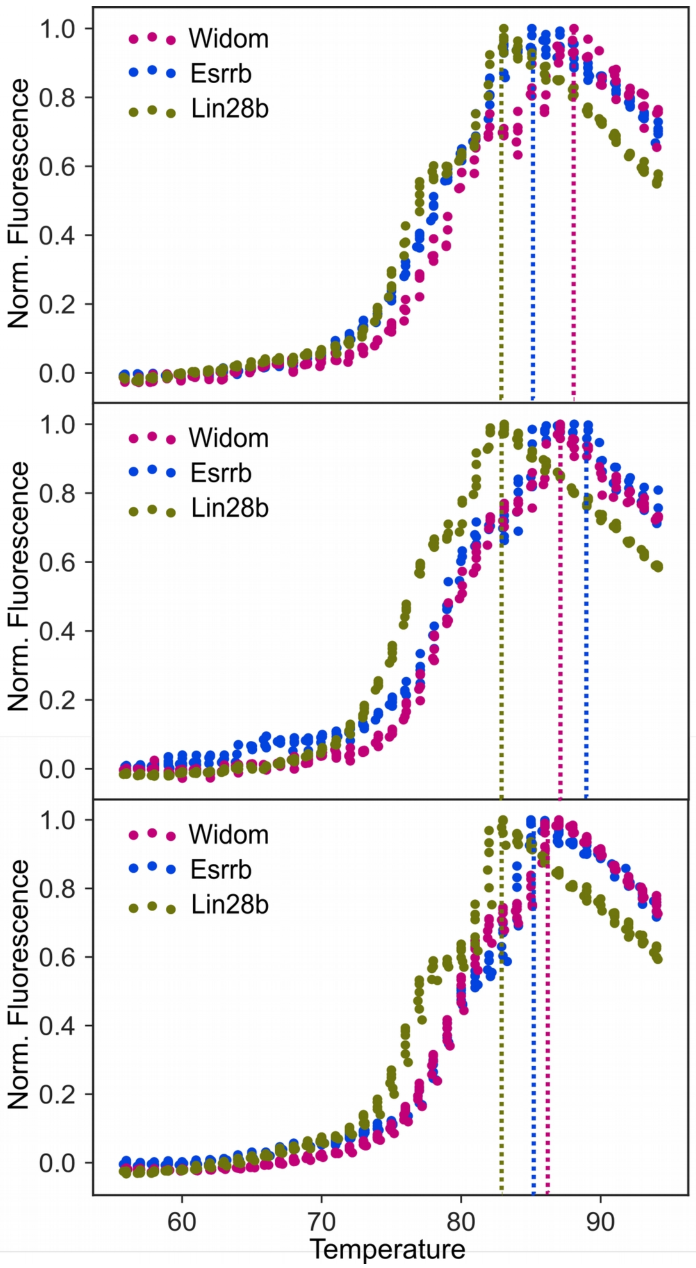


Huertas et al., Figure S1

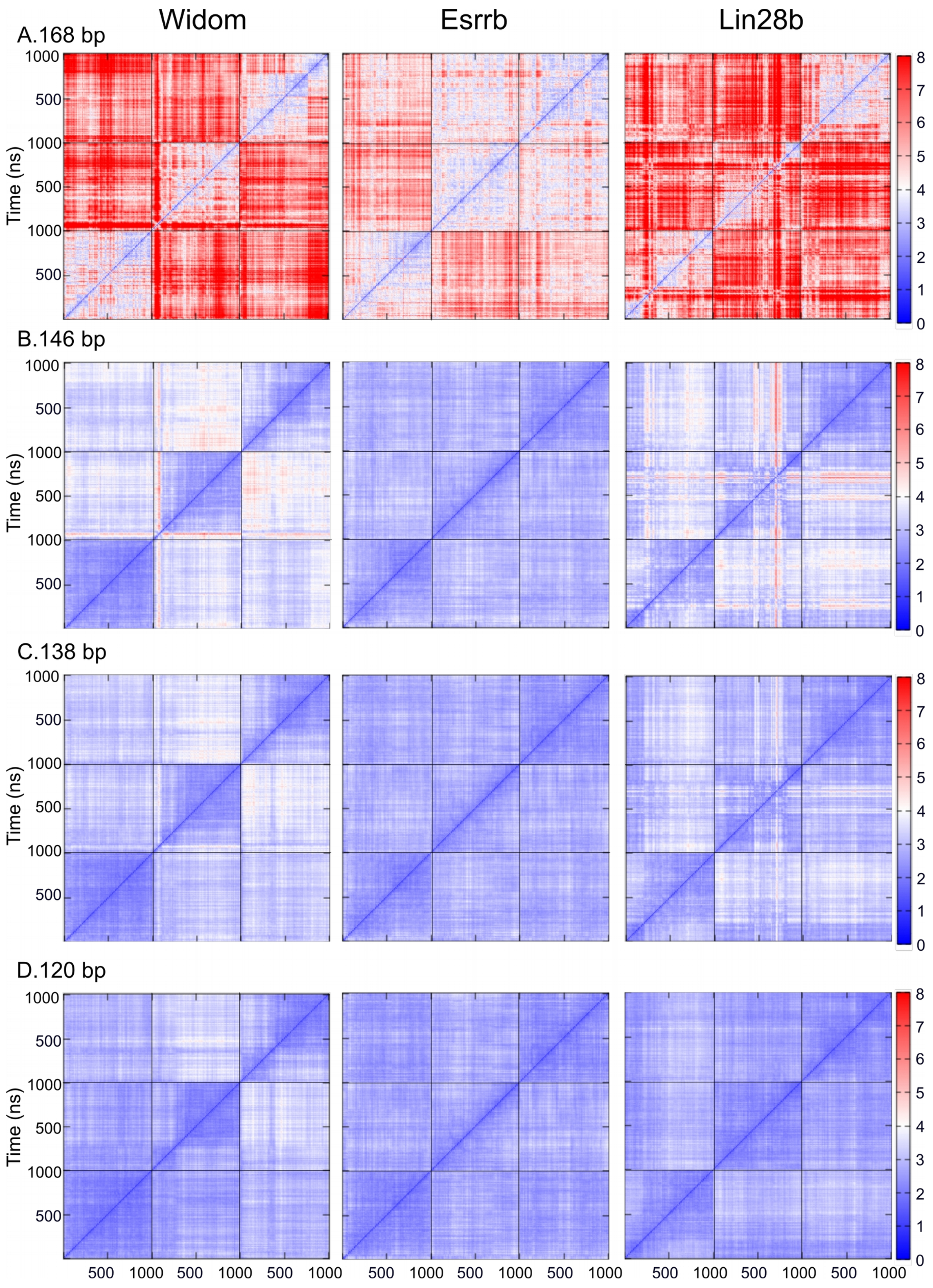


Huertas et al., Figure S2

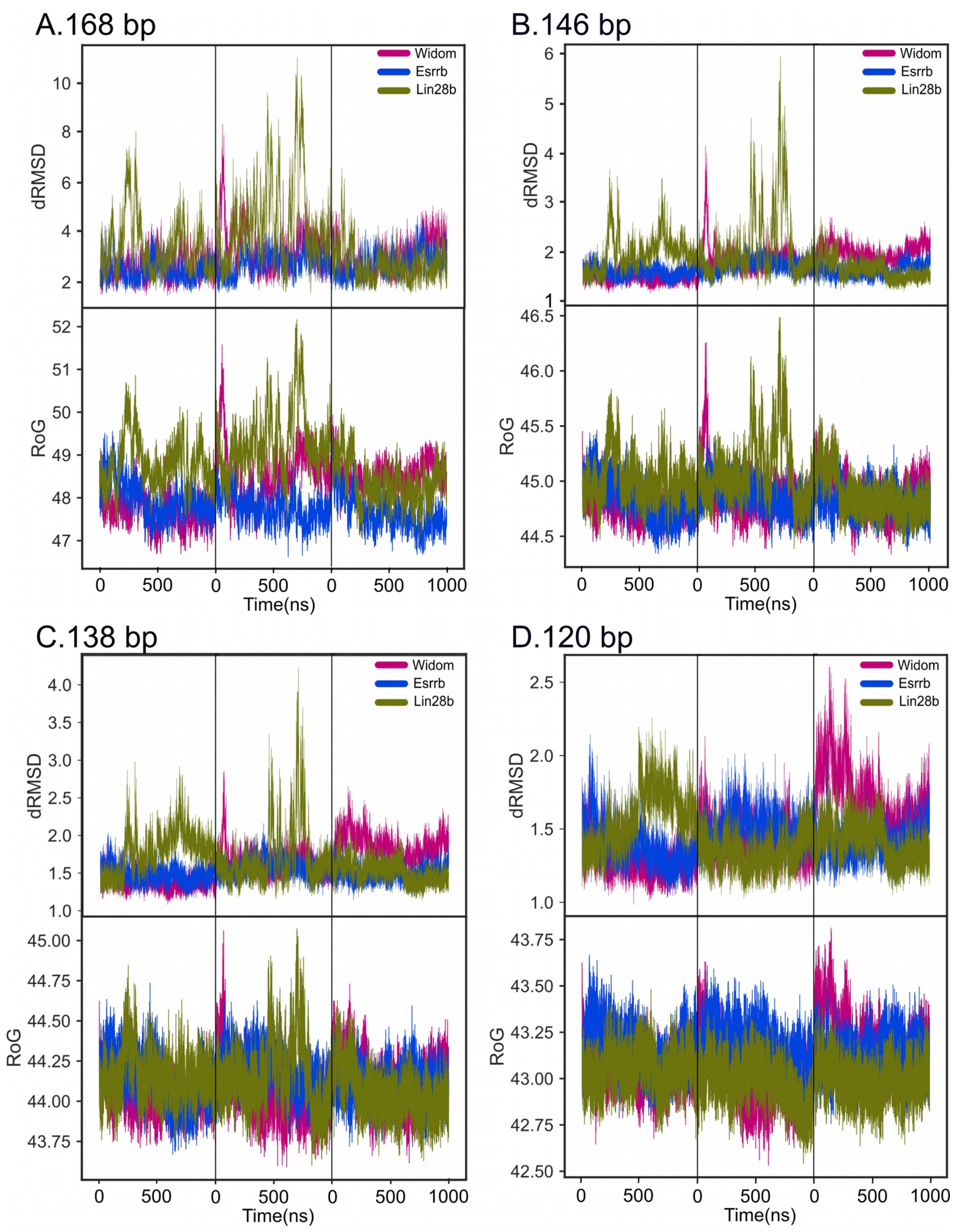


Huertas et al., Figure S3

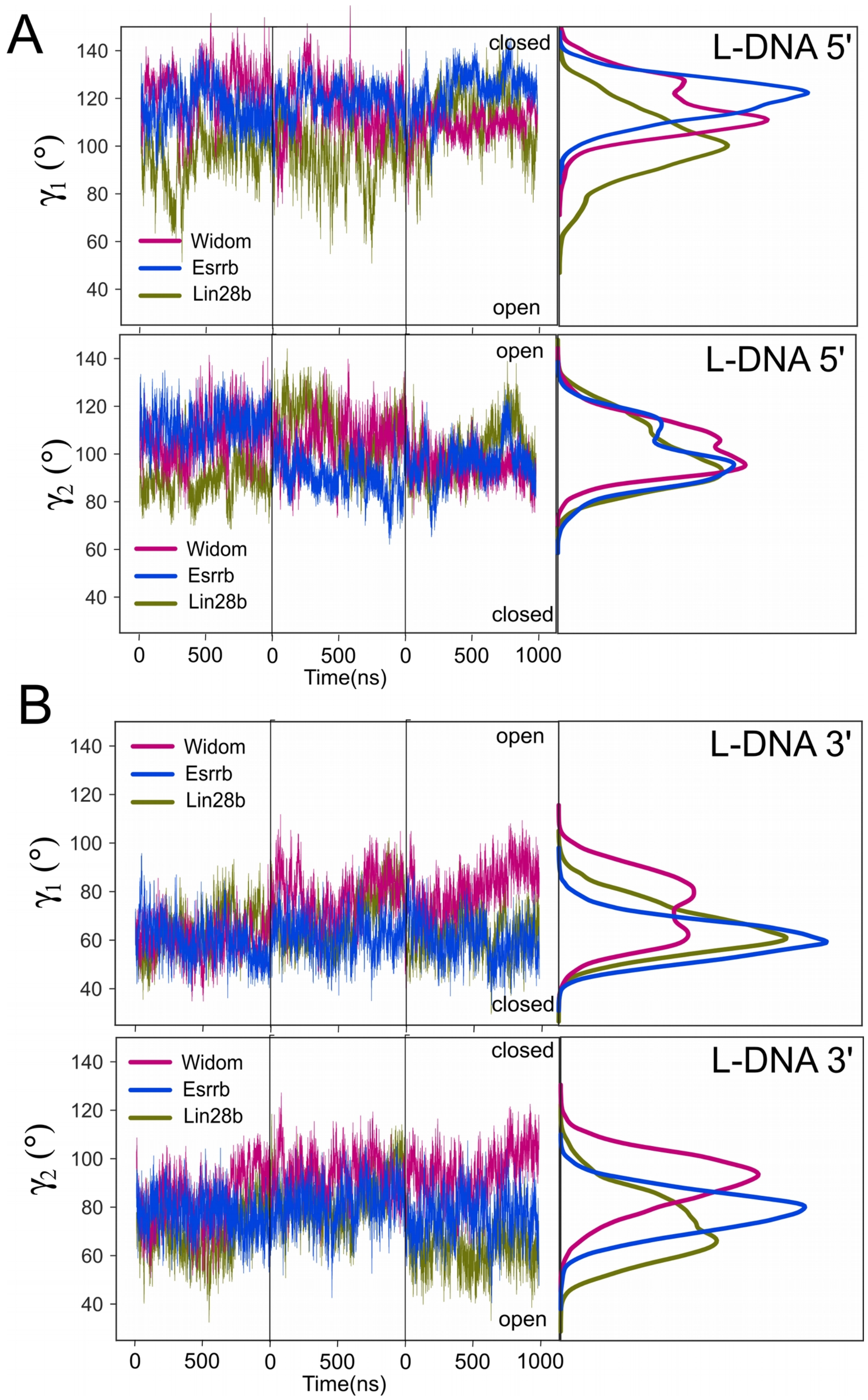


Huertas et al., Figure S4

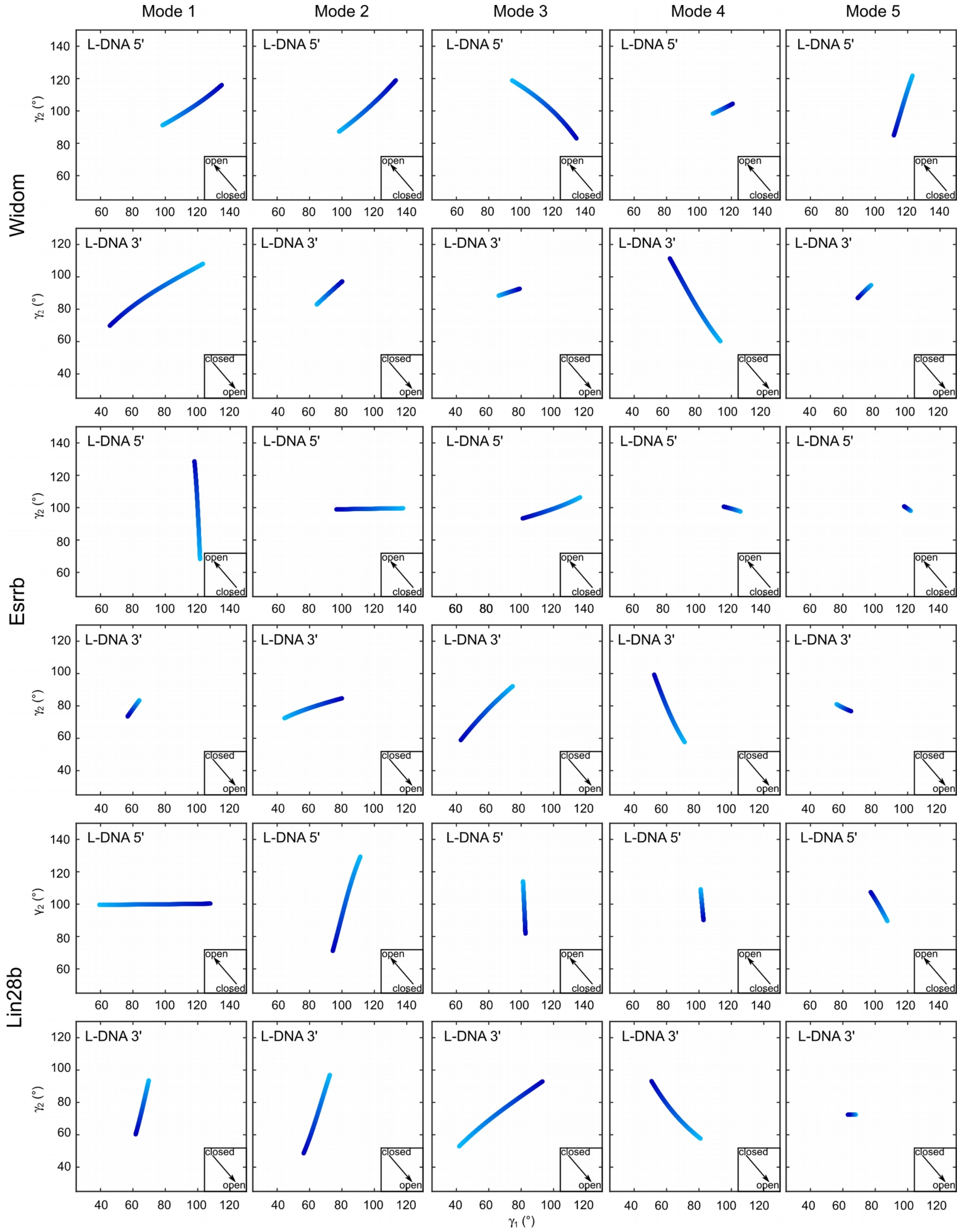


Huertas et al., Figure S5

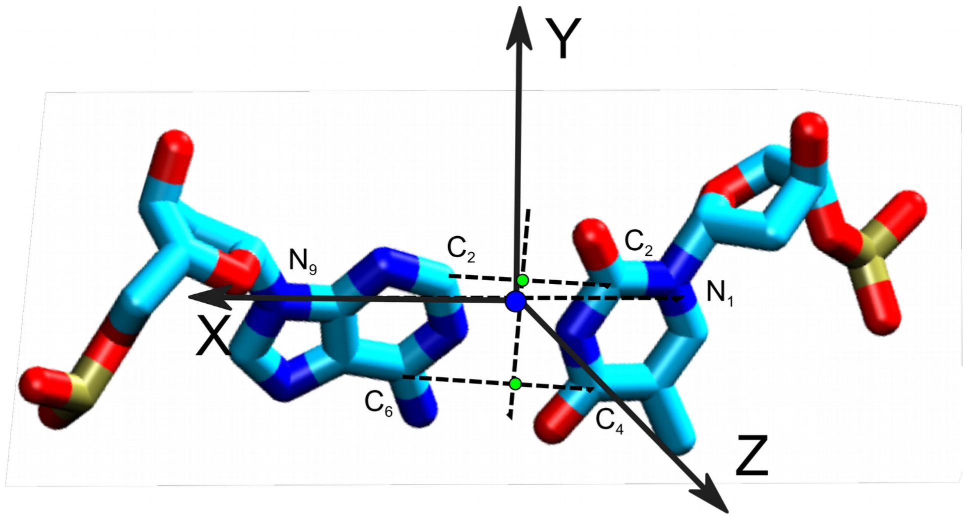


Huertas et al., Figure S6

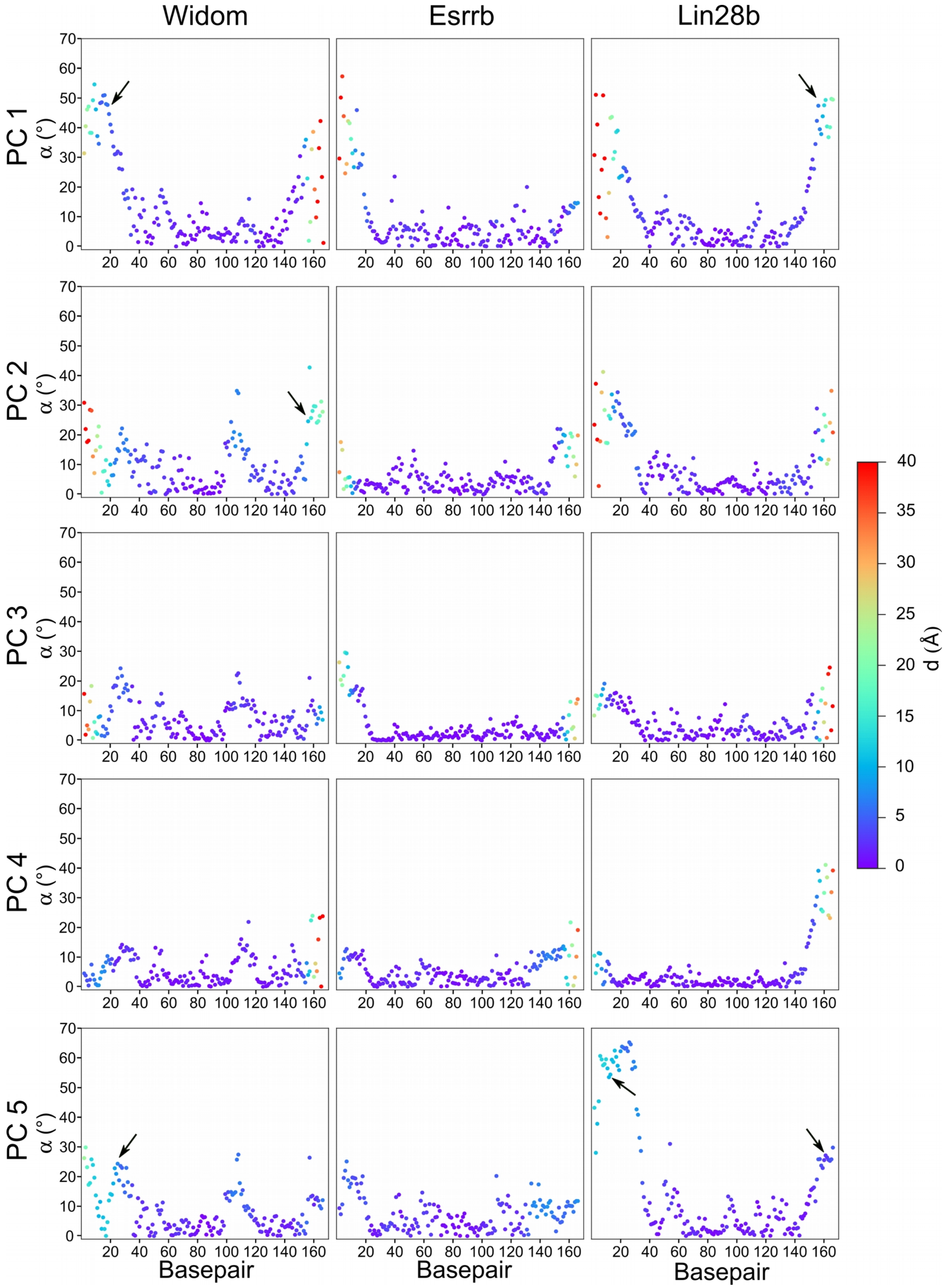


Huertas et al., Figure S7

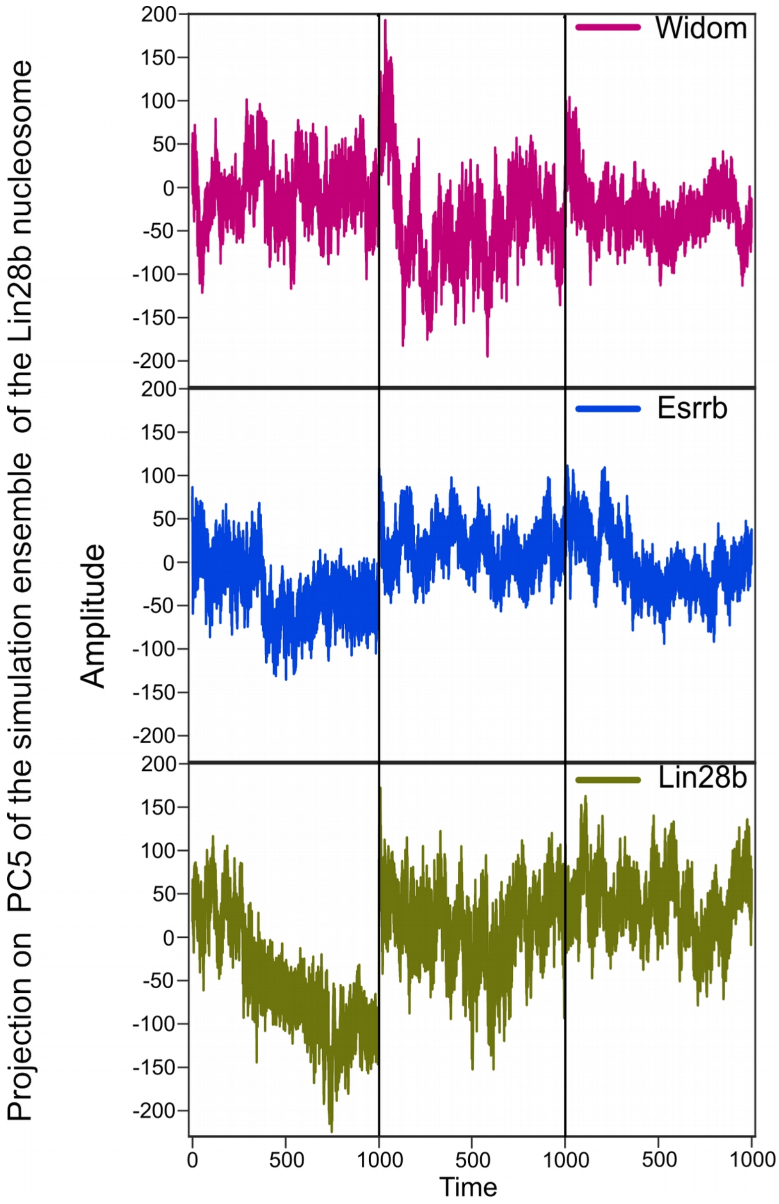


Huertas et al., Figure S8
